# Eelgrass (*Zostera* spp.) associated phytomyxids are host-specific congeneric parasites and predominant eukaryotes in the eelgrass rhizosphere on a global scale

**DOI:** 10.1101/2023.03.05.531089

**Authors:** Viktorie Kolátková, Megan Mooney, Kate Kelly, Elitsa Hineva, Ryan M. R. Gawryluk, Joel Elliott

## Abstract

Together with increasing environmental and anthropogenic pressures, pathogenic diseases are one of important factors contributing to the ongoing decline of seagrass meadows worldwide; yet the diversity and ecology of the microorganisms acknowledged as seagrass parasites remain critically understudied. Here we investigate phytomyxid parasites (Rhizaria: Endomyxa: Phytomyxea) of three different eelgrass (*Zostera* spp.) species found in the Northern hemisphere. We present molecular evidence that *Plasmodiophora bicaudata*, a long-recognized parasite of dwarf eelgrass taxa, is closely related to the novel phytomyxid recently discovered in root hairs of *Zostera marina*, and together they form a distinct clade within the order Phagomyxida, proposed here as *Feldmanniella* gen. nov. A full life cycle is systematically described in a phagomyxid representative for the first time, proving its conformity with the generalized phytomyxid life history, despite previous uncertainties. The presence of primary infection stages in nearly all collected eelgrass specimens, and subsequent analysis of amplicon sequences from a global *Z. marina* dataset, reveal phytomyxids to be ubiquitous and one of the predominant microeukaryotes associated with eelgrass roots on a global scale. Our discoveries challenge the current view of Phytomyxea as rare entities in seagrass meadows and suggest their generally low pathogenicity in natural ecosystems.

**Originality-Significance Statement:** This study addresses a group of microbial parasites critically understudied in the marine environment. It presents complex evidence that Phytomyxea - obligate intracellular biotrophs previously considered to be rare entities in the oceans, are in fact ubiquitous endobionts of seagrasses of the genus *Zostera* – foundation species and important primary producers in coastal areas worldwide. Our work represents a significant contribution to the fields of aquatic microbiology and seagrass ecology and is seminal to understanding the biology of Phytomyxea outside of terrestrial ecosystems.

## INTRODUCTION

Seagrasses of the genus *Zostera* (Alismatales: Zosteraceae), commonly referred to as ‘eelgrass’ and ‘dwarf eelgrass,’ are a group of true marine angiosperms with ribbon-like leaves known to form vast underwater meadows in estuaries and intertidal to subtidal zones worldwide (den Hartog 1970, Green and Short 2003). They are considered important ecosystem engineers, providing a variety of essential ecosystem services such as habitat and feeding ground for numerous animal assemblages (including commercially important fish and invertebrates), prevention of coastal erosion by reducing the hydrodynamic energy of waves, and stabilization of the seafloor sediment. (Bos et al. 2007, Kennedy et al. 2018, Surugiu et al. 2021, Meysick et al. 2022). With nine currently recognized species and a geographical range encompassing areas from the tropics to the subpolar regions, eelgrasses represent the most widely distributed and the second most diverse group of seagrasses altogether (Moore and Short 2007).

Despite their extensive area coverage and tolerance of environmental conditions such as different light regimes, temperatures and salinities, eelgrass meadows (like other seagrasses) have been experiencing a rapid decline in the past 100 years, triggering an immense global initiative for their conservation (Boström et al. 2014, Eriander et al. 2016, Orth et al. 2006, Orth et al. 2010, Valle et al. 2015, Xu et al. 2021). Large-scale eelgrass losses are commonly attributed to several biotic and abiotic factors, with extreme climatological events, anthropogenic disturbance and pathogenic diseases suspected to be the leading causes (Orth et al. 2006). However, the diversity and complex dynamics of eelgrass pathosystems are still only poorly understood (Sullivan et al. 2018). There are three groups of microbial organisms currently recognized as parasites of eelgrass: labyrinthulids, the causative agents of eelgrass wasting disease (Short et al. 1987, Sullivan et al. 2013); the *Phytophthora/Halophytophthora* species complex, which is thought to negatively affect eelgrass seed germination and viability (Govers et al. 2016); and phytomyxids, obligate intracellular biotrophs causing morphological deformations in the infected plants (Neuhauser et al. 2011). Interestingly, while research focused on the first two groups seems to be gaining momentum, seagrass-associated phytomyxids are perceived as rare, and are generally overlooked in contemporary eelgrass pathogen studies (*e*.*g*., Menning et al. 2020, 2021).

Infamous for infecting economically significant crops, Phytomyxea (SAR: Rhizaria: Endomyxa) is a monophyletic group of protists known to parasitize a wide range of hosts, including numerous terrestrial plants, seagrasses, brown and green algae, and oomycetes (Bulman and Neuhauser 2017). It currently comprises nearly 20 established genera accommodated in two major orders – the terrestrial/freshwater Plasmodiophorida and the marine Phagomyxida, and a recently described deep-branching marine clade ‘TAGIRI-5/Marinomyxa’ (Bulman and Neuhauser 2017, Hittorf et al. 2020, Kolátková et al. 2021). Similar to other endoparasites, the life cycle of phytomyxids is very complex and consists of numerous developmental stages typically taking place in three spatially separated environments: ‘primary’ sporangial infection of root hairs, ‘secondary’ sporogenic infection of cortical tissues, and sediment-dwelling resting spores (Bulman and Braselton 2014, Liu et al. 2020). This generalized scheme is, nevertheless, based mainly on the thoroughly studied representatives pathogenic to plants cultivated in agriculture, and may slightly differ in other phytomyxid taxa. In natural environments, the discoveries of novel phytomyxid species are sporadic and mostly tied to serendipitous observations of morphological changes in the host tissues. Typically, only a part of the life cycle is documented in the taxonomic descriptions, and a surprisingly large proportion of the recognized taxa has only been found on several occasions with old drawings being the only evidence of their existence (Karling et al. 1968). In the marine-derived Phagomyxida, a complete life cycle has not been observed in any of the species described so far, leading to prior speculations that some of the developmental stages may be missing in this lineage (Maier et al. 2000, Parodi et al. 2010, Bulman and Neuhauser 2017).

The first observation of phytomyxid parasites of the seagrass genus *Zostera* dates to 1938, when *Plasmodiophora bicaudata* was discovered in specimens of *Zostera noltei* from Tanoudert, Mauritania (Feldmann 1940). The parasite was reported to induce the formation of hypertrophies in the bases of the plant’s shoots and suppress the growth of internodes, giving the host a tufted ‘*Isoetes*-like’ appearance (Feldmann 1940, Feldmann 1956). In a subsequent herbarium study, den Hartog (1989) also confirmed the presence of *P. bicaudata* in *Zostera capensis* from South Africa, *Zostera muelleri* from Victoria (Australia), and *Zostera japonica* from British Columbia (Canada) and concluded that the parasite is rare, but widely distributed throughout populations of different dwarf eelgrass species. Hundreds of *Z. marina* specimens were also examined for *P. bicaudata* infection, but no signs of galls and unusual hypertrophies were found (den Hartog 1989). Despite its unknown pathogenic (and therefore ecological) potential, no further research on *P. bicaudata* was carried out, and secondary sporogenic plasmodia and resting spores with two characteristic appendages remain the only observed life stages of the organism so far (Feldmann 1940, Feldmann 1956). Furthermore, the lack of any molecular data has prevented verification of the morphometry-based taxonomic placement of the organism in the genus *Plasmodiophora*.

In 2015, a novel phytomyxid parasite was discovered in the roots of *Z. marina* in Puget Sound, WA (Elliott et al. 2019). This time, primary sporangial plasmodia and biflagellate zoospores were observed in the root hairs of infected plants, but no secondary infection or morphological deformations resembling *P. bicaudata* infection were detected. Phylogenetic analysis of the 18S rRNA gene placed the parasite within the phytomyxid order Phagomyxida comprising other marine representatives, with the closest relative being *Ostenfeldiella diplantherae* (86.9% nucleotide identity), a gall-forming parasite of the tropical seagrass *Halodule* (Ferdinandsen and Winge 1914, den Hartog 1965). However, no definite taxonomic treatment was given, since no genetic comparison to the previously described eelgrass parasite (*P. bicaudata*) could be performed. Since the observed developmental stages (primary sporangial plasmodia and biflagellate zoospores) conveniently complete the root hair component of the typical phytomyxid life cycle (as described above), we hypothesize that the organism investigated by Elliott et al. (2019) is either closely related to *P. bicaudata* or simply represents the so far overlooked life stages of the long-known phytomyxid species.

To resolve the identity of the newly discovered phytomyxid parasite infecting *Z. marina* and to understand its phylogenetic relationships with *P. bicaudata* and other seagrass-associated phytomyxid parasites, we examined specimens of several eelgrass species from the Northern hemisphere susceptible to phytomyxid infection and, for the first time, analyzed their parasites on a molecular level using rRNA-encoding phylogenetic markers. Moreover, we reviewed some previously published environmental sequencing amplicon datasets from eelgrass meadows around the world (Ettinger et al. 2021) to better understand the distribution of eelgrass-infecting phytomyxids on a global scale.

## EXPERIMENTAL PROCEDURES

### Seagrass tissue collection

Three different species of *Zostera* were collected for the purposes of this study. In the summer of 2018, a search for specimens of dwarf eelgrass, ***Zostera noltei***, infected with *Plasmodiophora bicaudata* was conducted at two distant localities in Europe (Table 1) at which the parasite’s presence had been previously reported in the scientific literature (den Hartog 1989, Hineva et al. 2017). In the mudflats of Zandkreek, Netherlands (Wadden Sea), a seagrass meadow was inspected during a low tide, and plants with a characteristic tufted appearance and bulbous shoot morphology (Fig. 1A) were harvested and stored in 70% ethanol for future examinations and DNA analyses. In Foros Bay, Bulgaria (Black Sea), plants with similar deformations were collected from depths of 0.2–0.3 m while snorkelling and were treated identically.

**Table 1.** List of seagrass specimens collected and examined in this study

| seagrass species | date of collection | locality | GPS | collection method | stored in | infection stage<sup>b</sup> |
| --- | --- | --- | --- | --- | --- | --- |
| *Zostera noltei*<sup>a</sup> | Jul 14, 2018 | Zandkreek, **Netherlands** | N 51.542335°<br>E 3.897222° | low tide | 70% EtOH | RHG + SG |
| *Zostera noltei*<sup>a</sup> | Sep 2, 2018 | Foros Bay, Burgas, **Bulgaria** | N 42.452311°<br>E 27.460873° | snorkelling (~0.3m depth) | 70% EtOH | RHG + SG |
| *Zostera japonica* | numerous,<br>2019-2021 | Dash Point, **WA, USA** | N 47.322412°<br>W 122.409332° | low tide | seawater | RHG, no SG found |
| *Zostera japonica* | May 28, 2021 | Tsawwassen Beach, Tsawwassen, **BC Canada** | N 49.013233°<br>W 123.119003° | low tide | seawater | RHG, no SG found |
| *Zostera japonica*<sup>a</sup> | Jun 25, 2021 | southern Roberts Bank, Delta, **BC Canada** | N 49.031271°<br>W 123.147920° | low tide | seawater | RHG, no SG found |
| *Zostera japonica*<sup>a</sup> | Aug 6, 2021 | southern Roberts Bank, Delta, **BC Canada** | N 49.031271°<br>W 123.147920° | low tide | 70% EtOH,<br>Karnovsky fixative | RHG + SG |
| *Zostera japonica*<sup>a</sup> | Sep 30, 2021 | Deep Cove, North Saanich, **BC Canada** | N 48.673176°<br>W 123.481448° | low tide | 70% EtOH,<br>Karnovsky fixative | RHG + SG |
| *Zostera japonica* | Sep 2021 | Birch Bay, **WA, USA** | N 48.930603°<br>W 122.754588° | low tide | 70% EtOH | RHG + SG |
| *Zostera marina* | numerous,<br>2019-2021 | Dash Point, **WA, USA** | N 47.322412°<br>W 122.409332° | low tide | seawater | RHG |
| *Zostera marina*<sup>a</sup> | Jun 23, 2021 | Cowichan Bay, **BC Canada** | N 48.742744°<br>W 123.624635° | low tide | seawater | RHG |
| *Zostera marina*<sup>a</sup> | Aug 6, 2021 | southern Roberts Bank, Delta, **BC Canada** | N 49.031271°<br>W 123.147920° | low tide | seawater | RHG |
| *Zostera marina* | Sep 17, 2021 | Plumper Cove, Keats Island, **BC Canada** | N 49.404057°<br>W 123.471639° | SCUBA (~3m depth) | seawater | RHG |
| *Zostera marina* | Nov 10, 2021 | Hope Bay, Pender Island, **BC Canada** | N 48.805845°<br>W 123.278205° | SCUBA (~3m depth) | seawater | RHG |

<sup>a</sup> Specimens from these collections were used for molecular analysis<sup>b</sup> RHG = root hair galls, SG = shoot galls

**Fig. 1.**
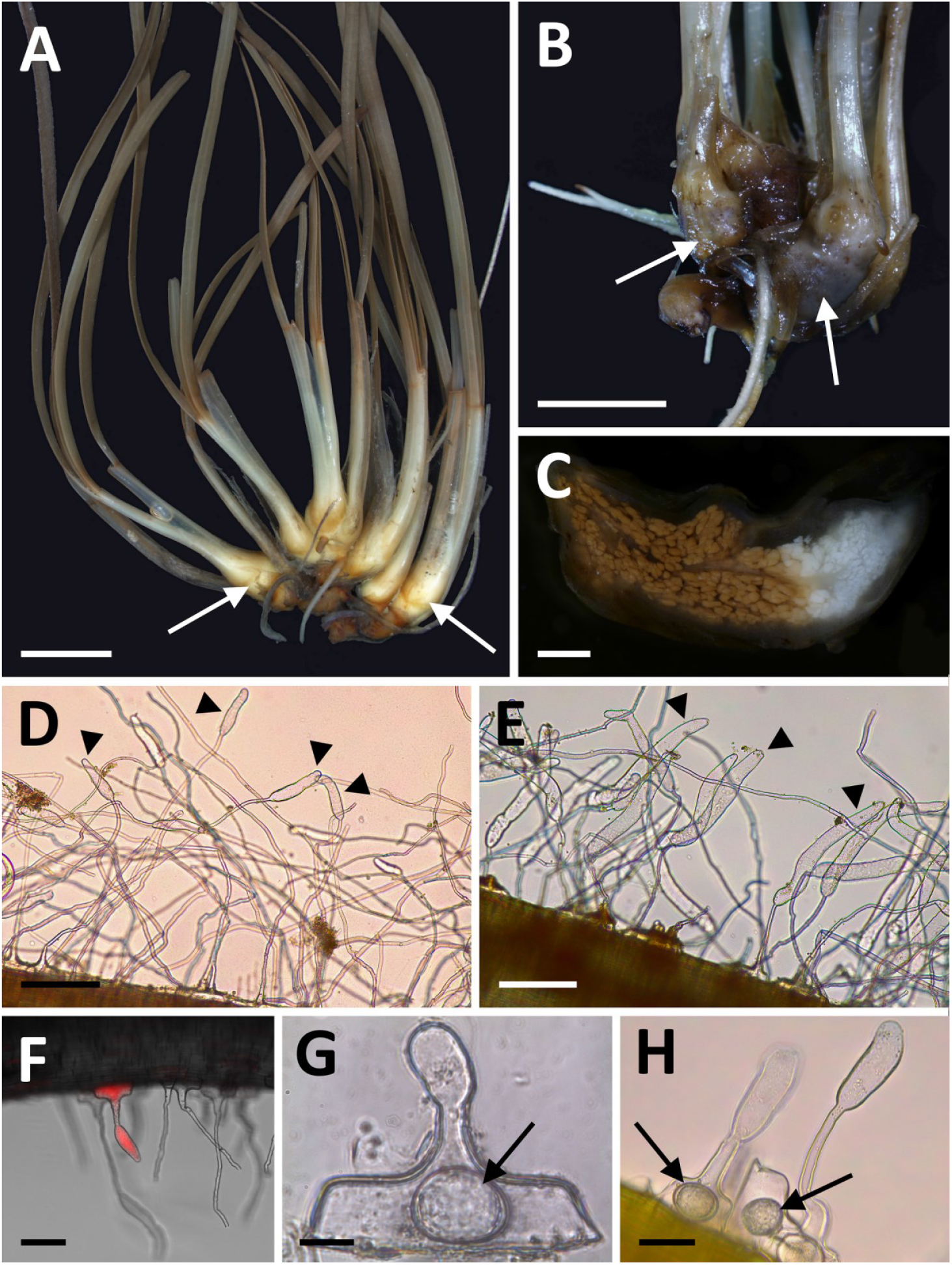
Morphology of seagrasses of the genus *Zostera* infested with phytomyxid parasites. (A) A specimen of *Z. noltei* heavily infected with *Feldmanniella bicaudata*. The secondary infection stages of the parasite induce the formation of galls in the bases of the seagrass shoots (arrows) and suppress the development of internodes, giving the plant an *Isoetes*-like appearance. (B) Shoot galls in a specimen of *Z. japonica* from the Salish Sea (Northeast Pacific) infected with *F. bicaudata*. (C) Cross section of a *Z. japonica* shoot gall revealing hypertrophied host cells filled with secondary sporogenic plasmodia (white coloration) and mature phytomyxid resting spores (brown coloration). (D) Root hair galls (arrowheads) in *Z. marina* containing primary developmental stages of *Feldmanniella radicapillae*. (E) Root hair galls (arrowheads) in *Z. japonica* containing primary developmental stages of *F. bicaudata*. (F) A root of *Z. marina* stained with Nile red showing the distribution of phytomyxid infection throughout the root hair. (G) Thick-walled ovoid structure of yet unknown function in the basal cell of an infected root hair of *Z. marina*. (H) Identical thick-walled ovoid structures in infected root hairs of *Z. japonica*. Scale bars: A, B = 5 mm; C = 1 mm; D, E = 200 µm; F = 100 µm; G = 20 µm; H = 50 µm.

In 2020–2021, numerous specimens of common eelgrass ***Zostera marina*** were collected during low tides from various sites throughout the Salish Sea, Northeast Pacific (Table 1) in search for root hair galls containing the novel phytomyxid parasite described by Elliott et al. (2019). Since the naked eye cannot detect the infection, random plants (including rhizomes and roots) were excavated from the surrounding sediment with a trowel, washed in ambient seawater, and stored in plastic zip-lock bags on ice in a cooler until transported to the laboratory. Within two hours after collection from the field, the rhizome and roots were carefully washed in artificial seawater (30 PSU, Instant Ocean®) to remove as much sediment as possible.

Alongside *Z. marina*, ***Zostera japonica*** (dwarf eelgrass invasive in Northeast Pacific) was also haphazardly collected at various localities in the Salish Sea (Table 1) and treated as described above to verify whether root hair galls are also formed in this species. Furthermore, following the finding of *Zostera japonica* being infected with *Plasmodiophora bicaudata* near Roberts Bank (BC, Canada) in 1985 (den Hartog 1989), several searches for the given symbiosis were carried out at this locality during low tides in May-August 2021. Plants with signs of infection (Fig. 1B) were placed in 70% ethanol for subsequent molecular analyses, and in Karnovsky fixative (Karnovsky 1965) for electron microscopy.

### Microscopy and measurements

#### Root hair galls

Individual roots of *Z. marina* and *Z. japonica* were cut off at their base with scissors and placed in a sonicator (FS 20, Fisher Scientific) filled with artificial seawater for approximately 10 sec to further remove sediment and other material adhering to the roots. The roots were stained with trypan blue or Nile red and examined using an inverted Leica compound microscope (DMIL LED) or an upright Leica compound microscope (DM 750) at 200–1000x magnification. Images were taken with a Leica EC3 camera and software. The length and width of randomly chosen root hair galls were measured using ImageJ software (Schneider et al. 2012). For scanning electron microscopy (SEM) root tissue was fixed in 4% glutaraldehyde in sterile seawater for two hours at 4°C and then dehydrated through an ethanol series (30, 50, 70, 90, 100%) and critical point drying using HMDS (hexamethyldisilazane) (Bray et al. 1993) Specimens were mounted on stubs and sputter coated with gold/palladium using a Denton Vacuum, Inc. sputter coater. The roots were then examined on a Hitachi S-3400N scanning electron microscope.

#### Secondary zoospores

A total of 50 cm of root tissue was put into Petri dishes with 10 ml of sterile seawater (0.45 µm filtered, 30 PSU) and incubated in the dark at 20°C for 24 hours. Zoospores released from the roots were examined at 400– 1000x magnification with a Leica inverted compound microscope and a Leica compound microscope. Measurements of zoospores were taken in ImageJ. Representative material was selected for further examination using scanning electron microscopy (SEM) and fixed in 4% glutaraldehyde in sterile seawater for two hours at 4°C. The fixative was gently poured out of the Petri dishes, and the zoospores left on the bottom were subsequently dehydrated through an EtOH series (30, 50, 70, 90, 100%) and critical point drying using HMDS (hexamethyldisilazane) (Bray et al. 1993). The bottoms of the Petri dishes were mounted on stubs and sputter coated with gold/palladium using a Denton Vacuum, Inc. sputter coater. The galls were then examined on a Hitachi S-3400N scanning electron microscope. Statistical analysis of all data was conducted using R (R Core Team 2015).

#### Sporogenic plasmodia and resting spores

Bulb-like hypertrophies at the bases of *Z. noltei* and *Z. japonica* shoots (hereinafter referred to as ‘shoot galls’) were inspected with an Olympus SZX9 stereomicroscope equipped with an Olympus DP72 camera. Several galls were sliced with a razor blade to reveal their internal arrangement (Fig. 1C). Plasmodial development was examined using transmission electron microscopy (TEM). *Z. japonica* gall tissue fixed in Karnovsky fixative (Karnovsky 1965) was post-fixed in 1.0% osmium tetroxide in 0.1 M cacodylate buffer (pH 7.2), dehydrated through an ascending ethanol series (15 mins at 30, 40, 50, 70, 80, 90, 95 and 2 × 100%), and embedded in EMbed 812 resin. Ultra-thin sections (80-90 nm) were cut using a Reichert Ultracut E ultramicrotome, mounted on 300 mesh copper grids, post-stained with uranyl acetate and lead citrate and viewed with a Jeol JEM 1400 TEM equipped with a Gatan SC-1000 digital camera. For SEM, part of the dehydrated tissue was not embedded in plastic, but critical-point dried with liquid CO_2_ in a Leica CPD300 critical point dryer, mounted on metal stubs, sputter-coated with gold and investigated using a Hitachi S-3500N scanning electron microscope with Quartz PCI software.

### Molecular methods

Roots of *Z. marina* and *Z. japonica* bearing root hair galls, as well as shoot galls found in *Z. japonica* and *Z. noltei* were used as source material for molecular analyses of phytomyxid DNA. For each host species, infected specimens from two different geographical locations were tested (Table 1). Plant tissue was homogenized in 1.5 ml microtubes with sterile plastic pestles and DNA was extracted using a DNeasy Plant Mini Kit (QIAGEN Sciences Inc., Germantown, MD) following the manufacturer’s instructions. For each sample, partial 18S and 28S rRNA genes and a ribosomal ITS region (containing the ITS1, 5.8S and ITS2 genes) were amplified with three different sets of phytomyxid-specific primers (Table 2). PCR reaction mixtures (25 μl) consisted of 12.5 μl of MegaFi Fidelity 2X PCR MasterMix (ABM Inc., Richmond, Canada), 1.5 μl of each primer (final concentration 0.6 μM), 1 μl of extracted DNA and 8.5 μl of ultrapure distilled H_2_O. To optimize the specificity of amplified fragments, touchdown PCR protocols were followed. For the amplification of the 18S and 28S rRNA genes, the protocol comprised an initial denaturation step at 98°C for 30 s, followed by 35 cycles of denaturation at 98°C for 10 s, primer annealing at 65°C for 30 s (in each cycle, annealing temperature was decreased by 1°C until 54°C was reached) and elongation at 72°C for 60s, and a final elongation step at 72°C for 5 mins. The same protocol was followed for the ITS region amplification, but the annealing temperature ranged from 62 to 56°C. PCR products were purified using the QIAquick PCR Purification Kit (QIAGEN Sciences Inc.), sequenced via ‘Sanger’ sequencing in both directions by Sequetech Corp. (Mountain View, CA) using various internal primers (Table 2), and manually assembled in BioEdit 7.0.5.3. (Hall 1999).

**Table 2.**
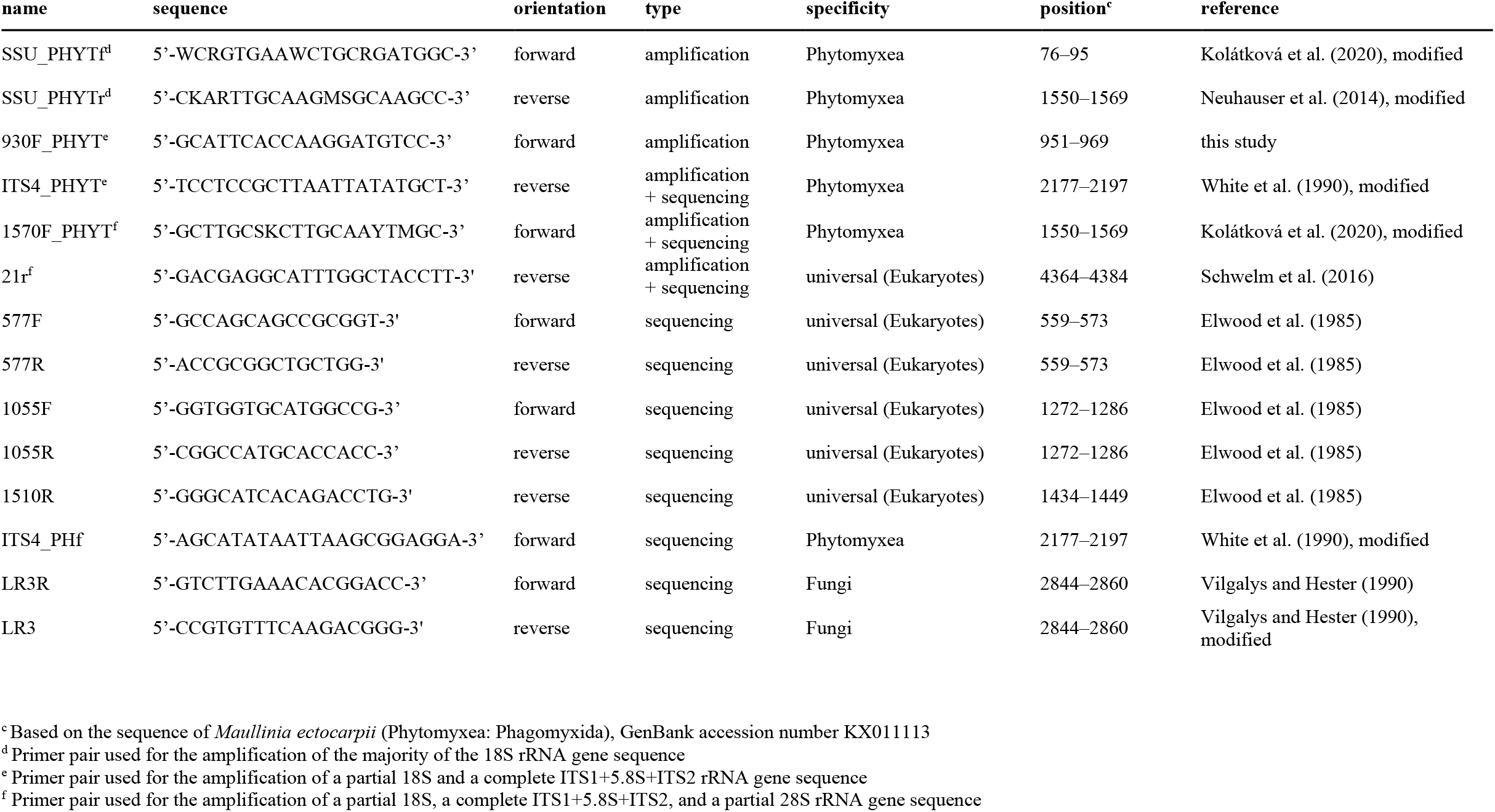
Oligonucleotide primer sequences used to amplify and sequence rRNA of examined phytomyxids

Approximately 4180 bp long consensus sequences covering a large part of the ribosomal operon were generated for the following phytomyxids: *Plasmodiophora bicaudata* in *Z. noltei* from Zandkreek, Netherlands; *P. bicaudata* in *Z. noltei* from Faros Bay, Black Sea; *P. bicaudata* in *Z. japonica* from Roberts Bank and North Saanich, BC (sequence identical between both locations and types of galls); and the novel phytomyxid in root hairs of *Z. marina* from Tsawwassen and Cowichan Bay, BC (sequence identical between both locations). The sequences were deposited in GenBank under the accession numbers OP137117-OP137120.

### Phylogenetic analyses

Phylogenetic analyses of the 18S and 28S rRNA genes were performed to determine the phylogenetic position of the examined protists within Phytomyxea. For the 18S rRNA gene phylogeny, a dataset containing the newly obtained sequences, a representative selection of 18S sequences of Phytomyxea, Endomyxa, and affiliated environmental sequences was generated and aligned using MAFFT (Katoh et al. 2002) on the MAFFT 7 server (https://mafft.cbrc.jp/alignment/server/) with the G-INS-i algorithm at default settings. The alignment was trimmed using trimAl v1.2 (Capella-Gutierrez et al. 2009) with ‘-gt 0.3 -st 0.001’ parameters to a final dataset containing 1598 aligned sites. Trees were reconstructed by maximum likelihood (ML) and Bayesian inference (BI) methods. The ML analysis was performed with IQ-TREE (Nguyen et al. 2015) using the TIM2+F+R4 model suggested as the best substitution model by ModelFinder (Kalyaanamoorthy et al. 2017), and 1000 nonparametric bootstrap replicates. The BI analysis was performed with Mr. Bayes v3.2.7 (Ronquist et al. 2012) using the GTR+I+ Γ model, with four rate categories for the gamma distribution. Four parallel runs of eight MCMCs were run for 20 million generations, with the first 5 million generations discarded as burn-in. The average standard deviation of split frequencies based on the remaining 15 million was < 0.01. Trees were sampled every 500th generation.

For the 28S rRNA gene phylogeny, the newly obtained sequences were aligned with a dataset of LSU rRNA sequences from Schwelm et al. (2016) and trimmed as described above, generating a final alignment containing 1811 sites. The ML analysis was performed with IQ-TREE using the TIM3+F+I+G4 model and 1000 nonparametric bootstrap replicates, and the Bayesian analysis was performed as described above.

### Analysis of global amplicon sequencing data

To better estimate the global distribution of phytomyxid parasites in eelgrass meadows, 18S amplicon sequencing data from a large study of the *Z. marina* mycobiome by Ettinger et al. (2021) were analyzed. Raw 2 × 300 bp paired-end Illumina V4 amplicon datasets, derived from 16 different geographical locations, were downloaded from the NCBI SRA repository (PRJNA667465). Adapter sequences 565F (CCAGCASCYGCGGTAATTCC) and 948R (ACTTTCGTTCTTGATYRA) were trimmed with CutAdapt v3.5 (Martin 2011). Only trimmed reads pairs in which both adapters were identified were retained. Trimmed reads were imported into QIIME2 v2021.11 (Bolyen et al. 2019). Amplicon sequence variants (ASVs) were inferred with DADA2 (Callahan et al. 2016), with forward and reverse reads truncated after 260 bp, based on visual inspection of quality scores. All samples with a total frequency of less than 5000 reads were discarded from the analysis.

ASVs corresponding to *Zostera*-associated phytomyxid parasites were identified by comparison to the sequences obtained in this study and the previously published 18S gene sequence (MG847140.1; Elliott et al. 2019). To facilitate treatment of two 99% identical ASVs from the phytomyxid parasite of *Z. marina* as a single entity in downstream analyses, while excluding the 97% identical ASV from the phytomyxid parasite of *Z. noltei/japonica*, ASVs were collapsed into operational taxonomic units (OTUs) at a 98% identity threshold. The abundance of the *Z. marina* parasite in each dataset was calculated as the percentage of its OTU counts, after excluding counts derived from *Z. marina* itself (Table S1 and S2).

## RESULTS

### Prevalence of phytomyxid infection in eelgrass meadows

The prevalence of shoot galls in the *Zostera noltei* and *Zostera japonica* meadows was generally very low (<1% of infected plants) and appeared to be seasonal. Infected specimens of *Zostera noltei* were found in summer/early autumn at both European localities (Table 1). During periodic searches for *Plasmodiophora bicaudata* in *Z. japonica* near Roberts Bank, BC, no shoot galls were found during sampling trips in May-early July, whereas in early August, specimens with visible hypertrophies were discovered readily. By September 2021, the presence of shoot galls was confirmed at all the *Z. japonica* sampling sites (Table 1). Infected plants appeared to be distributed in patches and at times could be recognized by visibly denser and shorter leaves compared to the neighbouring plants with unaltered morphology. In *Zostera marina*, no shoot galls were observed at all.

In contrast, root hair galls were present in nearly all specimens of *Z. marina* and *Z. japonica* collected in the Salish Sea throughout 2020 and 2021, regardless of location and season. Root hair galls were also observed in the roots of *Z. noltei* specimens collected in Europe but were not further examined due to their storage in ethanol. Despite the omnipresence of phytomyxid parasites, the seagrass meadows did not seem to be under distress. Seagrass uprooting resulting from phytomyxid infection, as previously proposed by den Hartog (1989), was not recorded at any of the sampling sites.

### Microscopy

#### Root hair galls

While the average diameter of root hairs in *Z. marina* (8.4+2.38 µm) and *Z. japonica* (8.35+2.01 µm) did not significantly differ (Wilcoxon, W= 9792, p=0.95), the average length and diameter of *Z. marina* root hair galls (Fig. 1D, length: 125+55 µm, diameter: 29.6+5.27 µm) were significantly smaller than those of *Z. japonica* (Fig. 1E, length: 249+76.8 µm, diameter: 36.3+4.61 µm) (length: t-test, t=16.06, df=295, p<0.0001, diameter: t-test, t=10.64, df=295, p<0.0001). Staining of roots with Nile red revealed that the infection is not limited to the gall itself, but parasitic biomass can be found throughout the entire root hair, including the basal part of the cell (Fig. 1F). Thick-walled structures located at the root hair bases previously reported by Elliott et al. (2019) were frequently observed in both seagrass species (Fig. 1G, H), but their identity/role remains unclear.

#### Secondary zoospores

In measurements from light microscopy images, the average cell length of *Z. marina* zoospores (5.77 + 0.38 µm) was significantly greater than those of *Z. japonica* (4.08 + 0.49 µm) (t-test, t=-12.66, df=43, p<0.0001). The long posterior flagella of the zoospores were clearly visible in light microscopy videos, but the shorter anterior flagella were difficult to observe (see Movie 1 and Movie 2). After fixation and dehydration for SEM, the cell body of the zoospores became smaller and more spherical in shape (Fig. 2A, B). In measurements from SEM images, average cell length of *Z. marina* zoospores (Fig. 2A; 3.02 + 0.21 µm) was significantly greater than those of *Z. japonica* (Fig. 2B; 2.57 + 0.29 µm) (t-test, t=-5.64, df=36, p<0.0001). The flagella were clearly visible in SEM images, and those of *Z. marina* zoospores were longer (anterior flagella = 4.54 + 0.56 µm, posterior = 14.44 + 1.45 µm), than for *Z. japonica* zoospores (anterior flagella = 2.45 + 0.96 mm, posterior = 11.4 + 1.46 mm). The anterior flagella were much shorter than the posterior flagella. The anterior flagella were on average 32% of the length of the posterior flagella in *Z. marina* and 18% of the length of the posterior flagella in *Z. japonica*. Fully developed secondary zoospores were observed leaving the root hair gall through small apertures in the gall’s cell wall (Fig. 2C, Movie 3). On several occasions, what appeared to be conjugating zoospores were captured both within (Movie 4) and outside of the root hair gall (Fig. 2D, E).

**Fig. 2.**
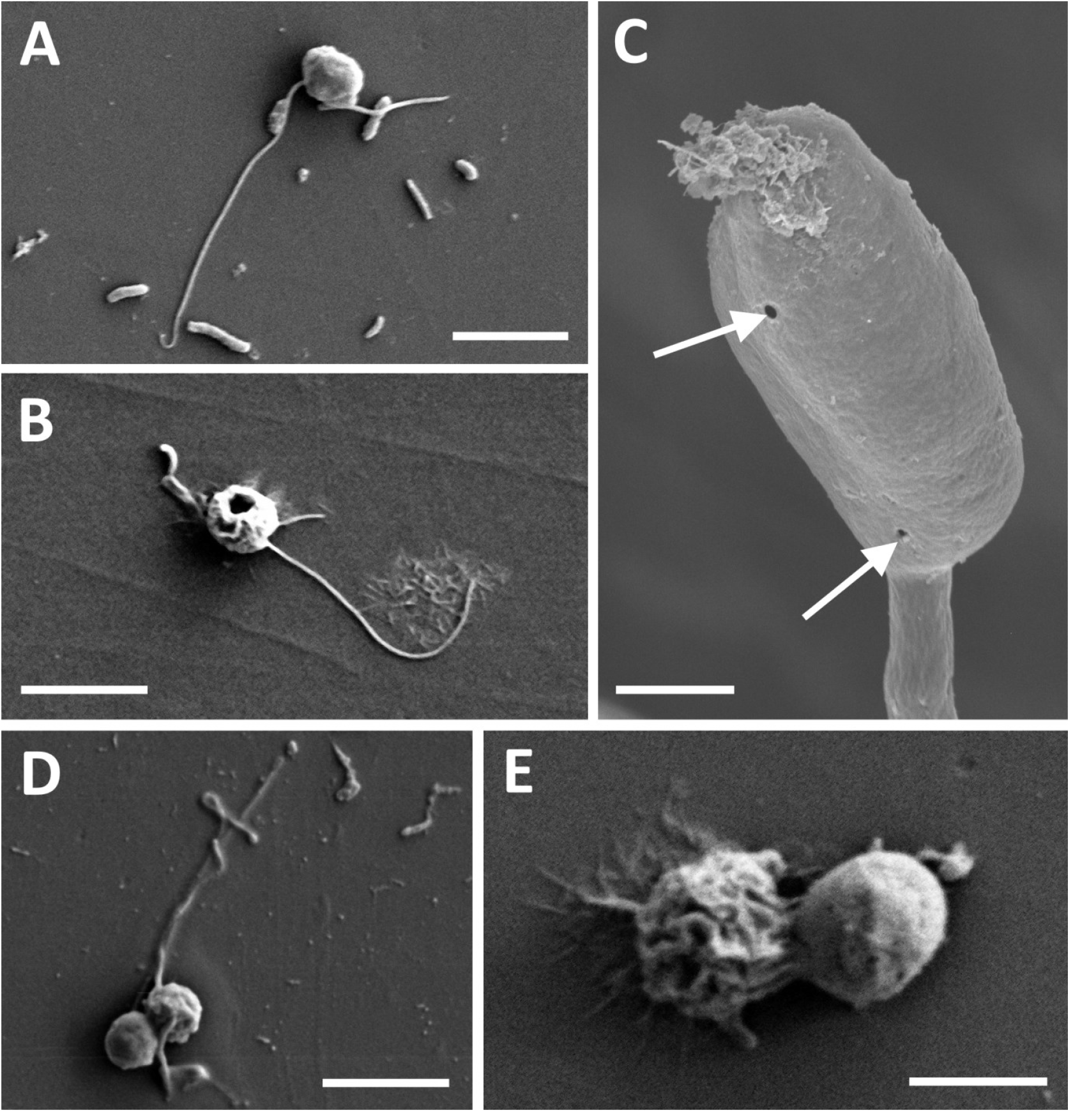
SEM micrographs of *Feldmanniella* spp. secondary zoospores. (A) Secondary zoospore of *F. radicapillae* released from a root hair gall in *Zostera marina*. (B) Secondary zoospore of *F. bicaudata* released from a root hair gall in *Zostera japonica*. (C) Apertures in a *Z. marina* root hair gall (arrows) used for the release of phytomyxid secondary zoospores (see Movie 3). (D, E) Apparent conjugation in secondary zoospores of *F. radicapillae*. Scale bars: A, B, D = 5 µm; C = 10 µm; E = 2 µm.

#### Sporogenic plasmodia and resting spores

The sliced shoot galls of *Z. japonica* and *Z. noltei* revealed greatly hypertrophied host cells within the cortical tissue of the seagrass leaves (Fig. 1C). The coloration of the host cells (white to dark brown) showed a distinct spatial distribution pattern of the parasite’s various developmental stages, with the youngest sporogenic plasmodia (white colour) located at the apical part of the gall and the mature resting spores located further down towards the rhizome (brown colour). Based on our TEM analysis, the secondary infection appeared to be induced by the presence of one or several uninucleate sporogenic plasmodia dwelling near the host cell wall (Fig. 3A). The growing plasmodia eventually filled out the host cell until only a degraded host nucleus remained, surrounded by a single large multinucleate plasmodium (Fig. 3B). At this phase, the plasmodium entered the ‘acaryotic’ transitional stage without visible nucleoli, during which meiosis occurs, and synaptonemal complexes can be observed in the nuclei (Fig. 3C). Cleavage into small uninucleate units followed, from which thick-walled spindle-shaped resting spores with the two characteristic appendages developed (Fig. 3D). SEM micrographs of the host cells further uncovered that these appendages are much longer than previously anticipated and maintain a unique intertwined spore arrangement (Fig. 3E).

**Fig. 3.**
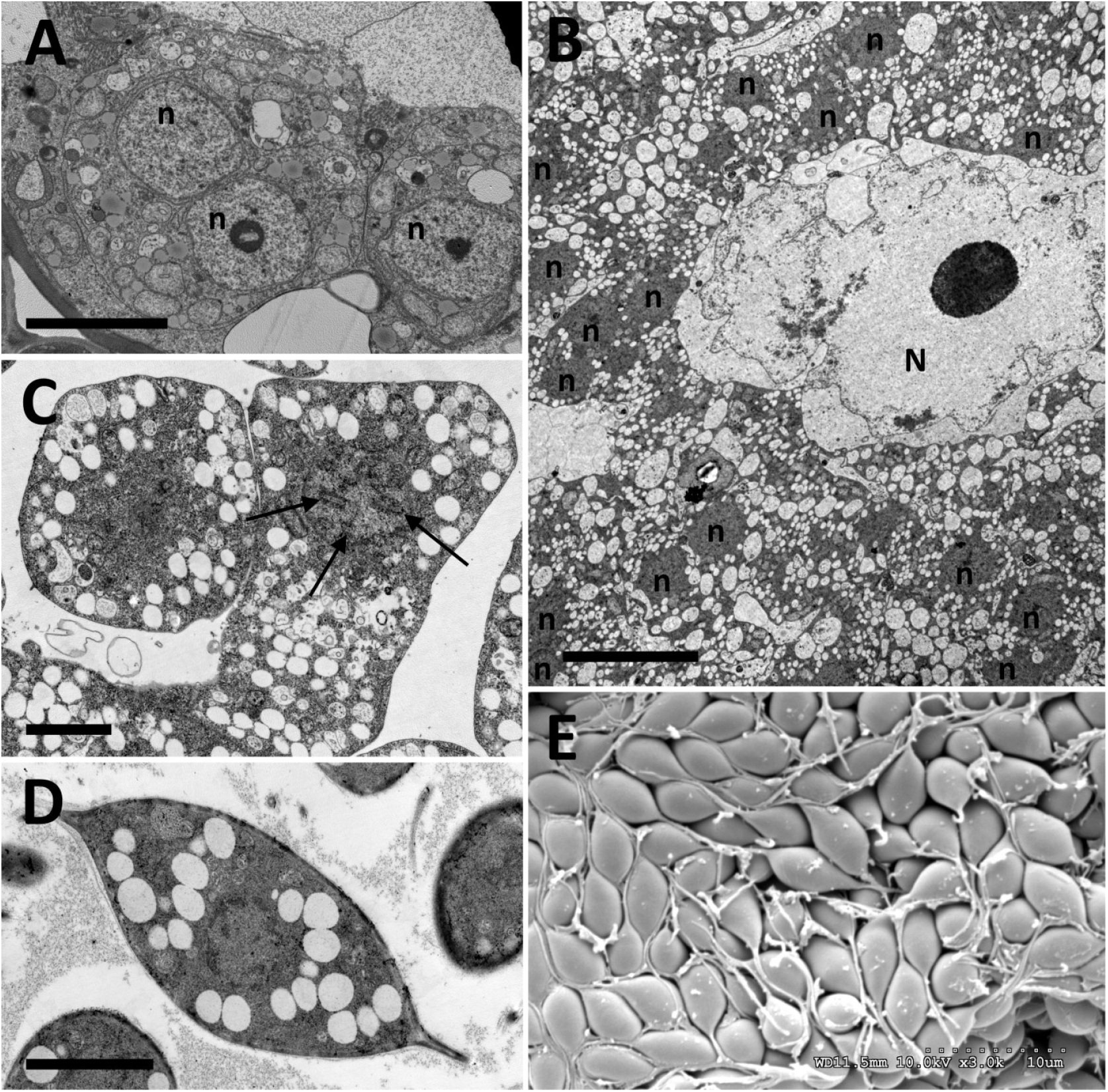
Secondary plasmodial development in *Feldmanniella bicaudata* (TEM and SEM). (A) Young uninucleate and binucleate (n) sporogenic plasmodia dwelling near the host cell wall. (B) Multinucleate (n) sporogenic plasmodium surrounding a degrading host nucleus (N). (C) Cleaving transitional sporogenic plasmodium in an ‘acaryotic’ stage. Parts of synaptonemal complexes are visible in the nuclei (arrows). (D) TEM micrograph of a fully developed resting spore with two characteristic appendages. (E) SEM micrograph of resting spore arrangement within the host cell. Scale bars: A = 5 µm; B = 10 µm; C, D = 2 µm; E = 10 µm.

### Phylogenetic analyses and genetic diversity

The phylogenetic trees of Phytomyxea inferred from the 18S and 28S gene sequences are shown in Fig. 4 and Fig. 5 respectively. In the 18S tree, all three major phytomyxid lineages (*i*.*e*., Plasmodiophorida, Phagomyxida and Marinomyxa/TAGIRI-5) were recovered with maximum support (bootstrap support ‘BS’ 100, Bayesian posterior probability ‘BPP’ 1). The newly obtained sequences of eelgrass-associated phytomyxids formed a clearly defined clade within Phagomyxida and were closely related to another seagrass parasite *Ostenfeldiella diplantherae* and the ‘TAGIRI-4’ environmental sequences (BS 86, BPP 1). While the support for this clade was poor when the long branching *O. diplantherae* was included in the analysis, it was robust (BS 100, BPP 1) when the *O. diplantherae* sequence was removed (Fig. S1). Our 18S sequence of the parasite forming root hair galls in *Z. marina* was nearly identical (99.5% identity) with the previously published sequence MG847140.1 from Puget Sound, WA (Elliott et al. 2019) and together they formed a distinct, well-supported (BS 97, BPP 1) branch within the novel clade. 18S sequences of the parasites found in *Z. japonica* and *Z. noltei* were also highly similar to each other (>99.5% identity) but their exact phylogenetic positioning within the clade was not definitively resolved by the phylogenetic analyses.

**Fig. 4.**
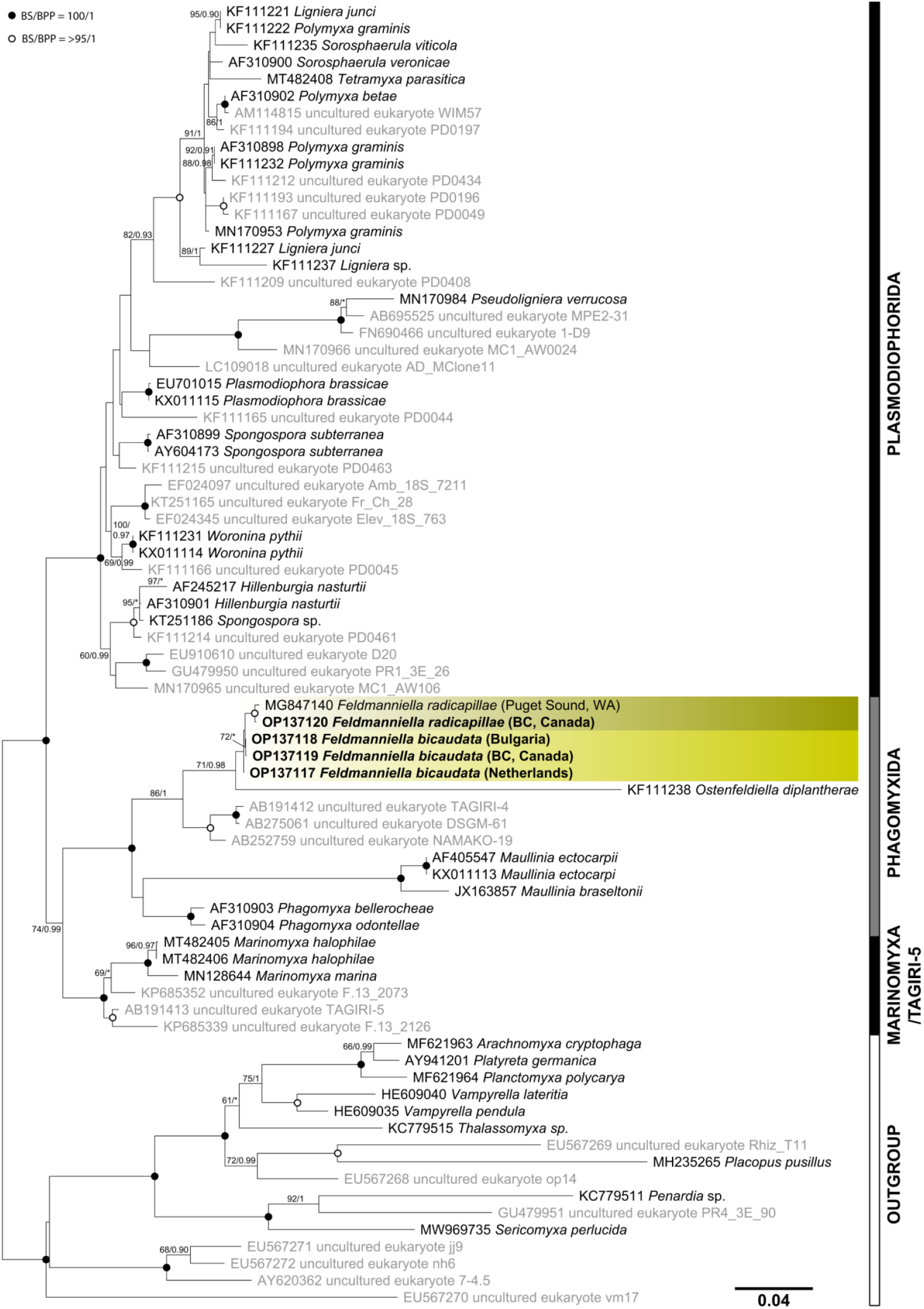
Phylogenetic tree of Phytomyxea based on maximum likelihood phylogenetic analysis of the 18S rRNA gene (1598 sites; 77 sequences). Yellow-green boxes highlight representatives of the newly established genus *Feldmanniella*. Sequences derived from isolates are in black, newly determined sequences of *Feldmanniella* spp. are in bold letters. Values at branches represent statistical support in bootstrap values (IQ-TREE)/ posterior probabilities (MrBayes); support values below 60/90 are not shown or are represented by an asterisk.

**Fig. 5.**
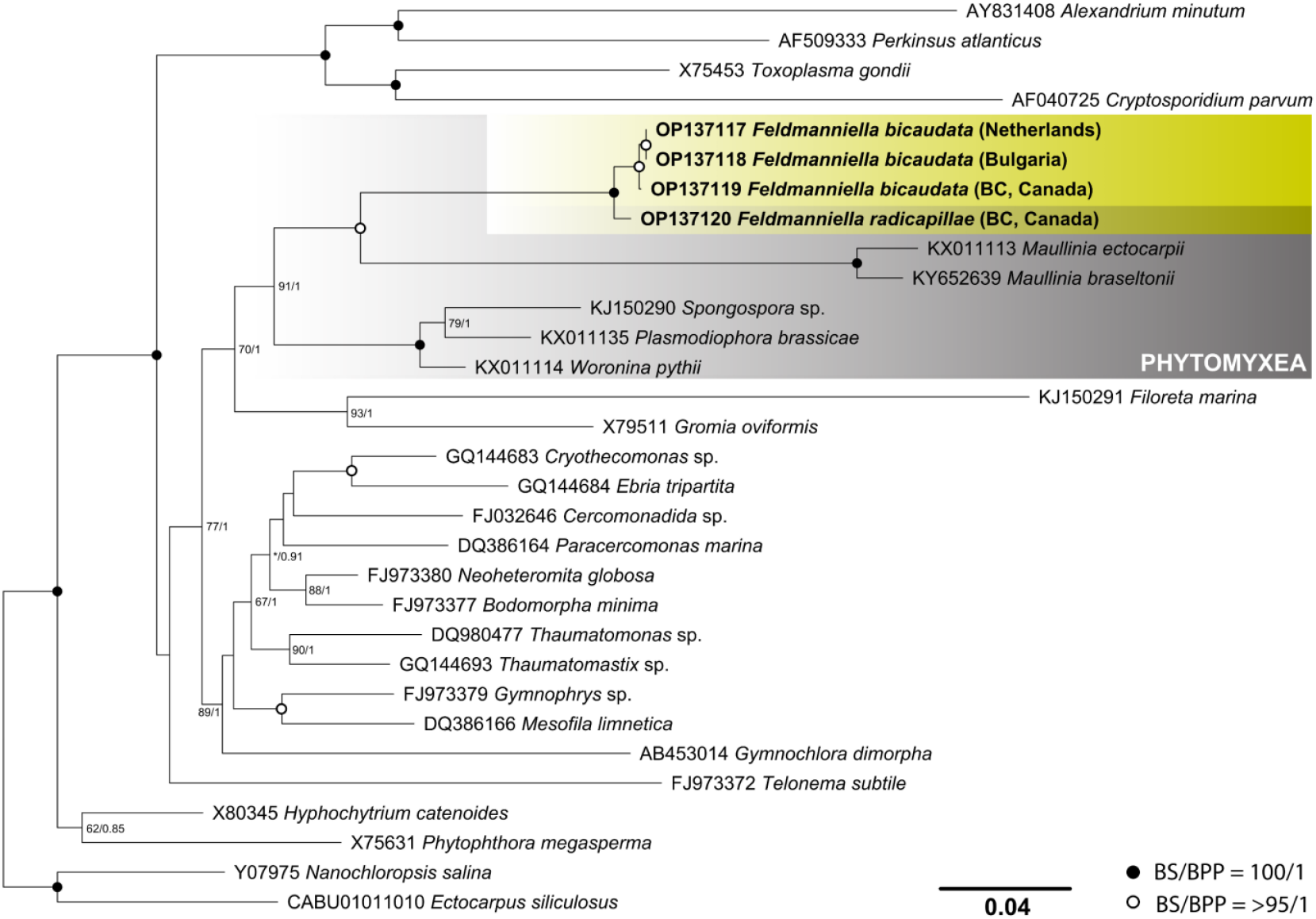
28S rRNA phylogenetic tree constructed by maximum likelihood analysis in IQ-TREE (1811 sites; 31 sequences chosen based on Schwelm et al. (2016)). Sequences of Phytomyxea are shaded in grey, yellow-green boxes highlight representatives of the newly established genus *Feldmanniella*. Sequences newly determined in this study are in bold letters. Values at branches represent statistical support in bootstrap values (IQ-TREE)/ posterior probabilities (MrBayes); support values below 60/80 are not shown or are represented by an asterisk.

The 28S gene tree further underlined these findings, with eelgrass-associated phytomyxids forming a robustly supported clade closely related to sequences of *Maullinia* spp. (Phagomyxida), parasites of brown algae. The newly determined sequences of the phytomyxids infecting dwarf eelgrass species (*i*.*e*., *Z. noltei* and *Z. japonica*) clustered together, whereas the phytomyxid from the roots of *Z. marina* was once more recovered as a distinct branch within the clade.

### Analysis of global amplicon sequencing data

The OTU of the phytomyxid parasite associated with *Z. marina* was found in datasets from *Z. marina* meadows from all 16 localities situated across the Northern hemisphere (Fig. 6). More specifically, it was detected in 85/212 analysed leaf samples, 140/166 analysed root samples and 101/164 analysed sediment samples (Table S2). The phytomyxid was highly abundant primarily in the root samples, where it often represented one of the most abundant OTUs altogether. In root samples from southern California, southern Japan, and North Carolina, the parasite’s OTU reached the second highest number of read counts, right after the OTU representing *Z. marina*. After the exclusion of *Z. marina* OTU from the dataset, the parasite’s OTU exceeded 10% of the root derived read counts at 5 out of the 16 localities, with the maximum value reaching 35.28% of read counts in southern California. At the geographical locations where dwarf eelgrass species can be commonly found (such as Croatia, Portugal and Washington) the OTU representing *P. bicaudata* was also detected in numerous samples.

**Fig. 6.**
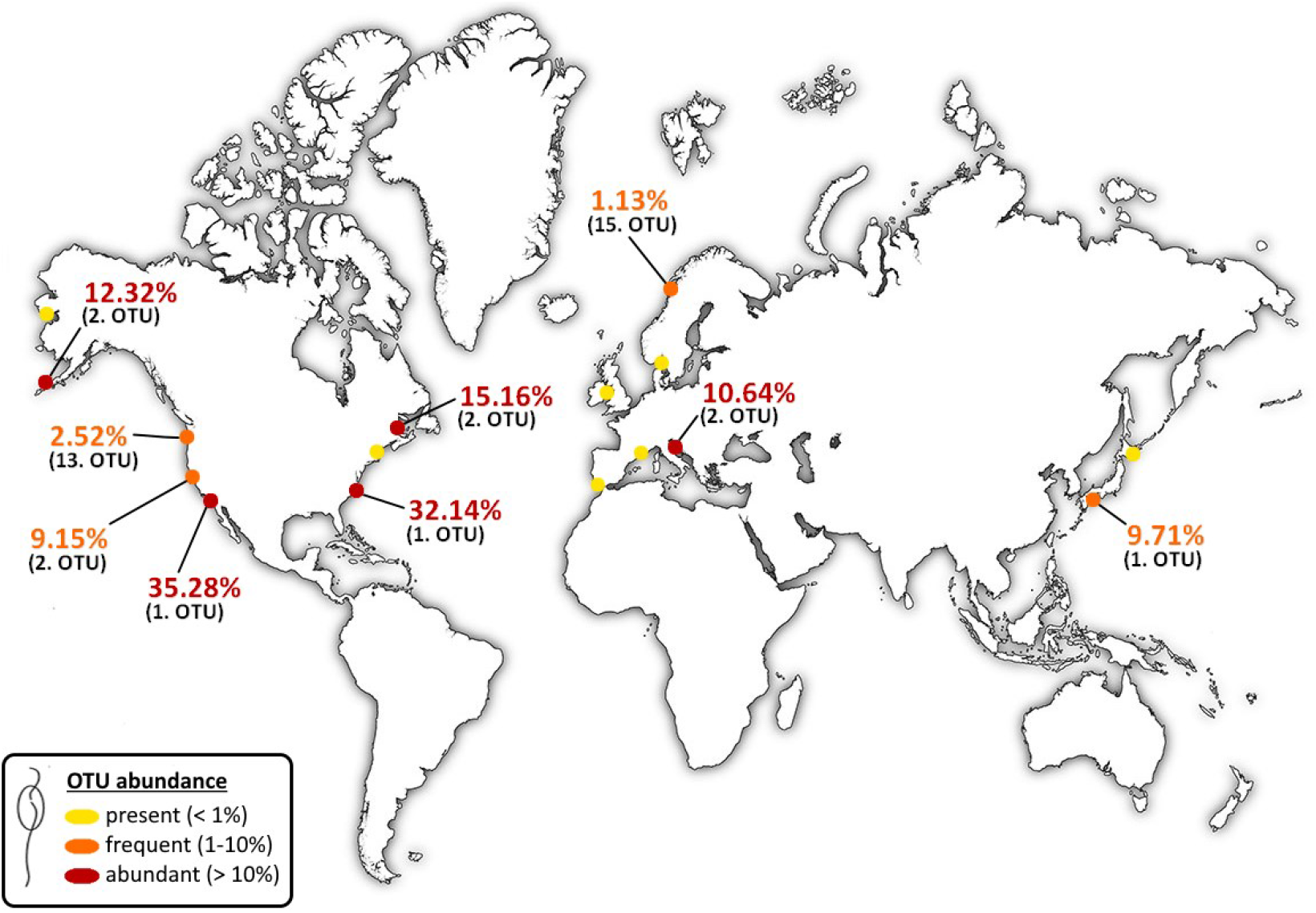
World map depicting 16 localities (circles) where specimens of *Zostera marina* leaves, roots and surrounding sediments were collected for an amplicon sequencing study conducted by Ettinger et al. (2021). Portrayed percentage values represent relative numbers of read counts belonging to the *Feldmanniella radicapillae* OTU in collected root samples at a given locality after removing *Z. marina* OTU counts from the dataset (values below 1% are not shown). Numbers in parentheses show the ranking of *F. radicapillae* OTU based on the total number of read counts from all root samples at a given locality; *Z. marina* OTU is not included in the ranking as it had the highest number of read counts at all sites.

### Taxonomic Summary

### Endomyxa: Phytomyxea: Phagomyxida

### *Feldmanniella* Kolátková, Mooney, Kelly, Hineva, Gawryluk et Elliott, gen. nov. Type species: *Plasmodiophora bicaudata* Feldmann, 1940

#### Included species

*Feldmanniella bicaudata* (Feldmann, 1940) comb. nov., *Feldmanniella radicapillae* (Kolátková, Mooney, Kelly, Hineva, Gawryluk et Elliott, 2022) sp. nov.

#### Etymology

The name refers to Jean Feldmann (1905–1978), an exceptional aquatic botanist and the discoverer of the first representative of this genus; and his wife Geneviève (1910–1994) who first described the cytological development of the respective organism.

#### Description

Ubiquitous, globally distributed endoparasites of seagrasses of the genus *Zostera* (Alismatales: Zosteraceae). Primary multinucleate plasmodia develop into zoosporangia within the host’s root hairs where they induce formation of cylindrical root hair galls. Released secondary zoospores are of an oval to pyriform shape and have two smooth heterokont flagella. Secondary multinucleate plasmodia develop in the cortex of the basal parts of the seagrass shoots, undergo meiosis and cleave into unicellular units, which eventually mature into yellow, spindle-shaped resting spores with two rigid appendages.

### *bFeldmanniella bicaudata* (Feldmann) Kolátková, Mooney, Kelly, Hineva, Gawryluk et Elliott, com. nov

#### Synonyms

*Plasmodiophora bicaudata* Feldmann, J., Bulletin de la Société d’Histoire Naturelle de l’Afrique du Nord 31: 171–177 (1940)

#### Original description

“Myxamoebae ignotae. Plasmodia plurinucleata, nucleis 3-4 µm longis, intra cellulas corticales eximie inflatas apicis caulis *Zosterae nanae* Roth. parasitica. Sporae singulae, totam cellulam hospitis replentes, luteolae vel in cumulo fuscescentes (in speciminibus in alcoholo servatis), ovoideae, 7 µm longae 3-3.5 µm latae utrinque in setulam acutam productae” (Feldmann 1940).

#### Note

In addition to Feldmann’s observation (Feldmann 1940), *F. bicaudata* was observed in herbarium specimens of *Zostera capensis, Zostera muelleri, Zostera japonica* and possibly also *Zostera capricorni* (den Hartog 1989). In cytological study of infected specimens of *Zostera noltei* from southern France (Feldmann 1956), the secondary plasmodia were interpreted as diploid with meiosis occurring prior to cleavage into resting spores. The fusion of zoospores or amoebae was hypothesized due to lack of evidence of karyogamy in the plasmodium (Feldmann 1956). Furthermore, seasonality in the life cycle was observed with meiosis occurring in August and September, but not earlier in the year (Feldmann 1956).

#### Addition to the description

Root hair galls containing primary sporangial plasmodia are ubiquitous and can be found year-round. In *Z. japonica*, the root hair galls are 249+76.8 µm long and 36.3+4.61 µm wide. Secondary zoospores are 3.5-4.5 µm long with posterior flagellum measuring 10-13 µm and anterior flagellum ∼2.5 µm.

### *Feldmanniella radicapillae* (Feldmann) Kolátková, Mooney, Kelly, Hineva, Gawryluk et Elliott, sp. nov

#### Description

Parasite of the seagrass *Zostera marina*. Root hair galls containing primary sporangial plasmodia are 125+55 µm long and 29.6+5.27 wide. They are ubiquitous and can be found year-round. Secondary zoospores are 5.4-6.1 µm large with posterior flagellum measuring 13-16 µm and anterior flagellum 4-5 µm. Secondary sporogenic plasmodia and resting spores have not been observed.

#### Etymology

The name indicates that the parasite was discovered in the root hairs of its host *Z. marina* (Elliott et al. 2019).

## DISCUSSION

Our molecular analyses of phytomyxid parasites from three different species of seagrasses of the genus *Zostera* revealed that the newly discovered phytomyxid from Puget Sound, WA (Elliott et al. 2019) and *Feldmanniella* (formerly *Plasmodiophora*) *bicaudata* (Feldmann 1940) are closely related, yet morphologically distinguishable organisms, which together form a novel clade within the order Phagomyxida, described here as *Feldmanniella* gen. nov. In agreement with conclusions made by den Hartog (1989), *F. bicaudata* was found only to parasitize dwarf eelgrass species (*i*.*e*., *Zostera noltei* and *Zostera japonica*) whereas the novel *Feldmanniella radicapillae* is a host-specific parasite of *Zostera marina*. This was true of all seagrass specimens obtained in the Salish Sea, including the ones from mixed meadows of *Z. japonica* and *Z. marina* near Roberts Bank, BC and Dash Point, WA (see Table 1), where the two seagrasses grew concurrently. The strong host specificity and close phylogenetic relations of the examined protists correspond with an identical pattern previously observed in *Marinomyxa* spp. – parasites of the (sub)tropical seagrass genus *Halophila* (Kolátková et al. 2021), and further support the hypothesis of a long coevolutionary history between phytomyxids and their seagrass hosts (Kolátková et al. 2021).

Despite prior speculations that the life history of Phagomyxida may be different and possibly less complex than that of Plasmodiophorida (Maier et al. 2000, Parodi et al. 2010, Bulman and Neuhauser 2017), all the developmental stages characteristic for the generalized plasmodiophorid life cycle (Bulman and Neuhauser 2017) are definitely present in *Feldmanniella*. The observations given here and in the cytological study conducted by Feldmann (1956) are highly congruent with the life cycle of *Plasmodiophora brassicae* recently refined by Liu et al. (2020), although several life stages need further investigation to be fully understood. The frequent occurrence of the ovoid thick-walled structures (possibly mature primary zoosporangia) in the bases of infected root hairs (see Fig. 1) suggests that they play a significant part in the infection process, which we have not uncovered yet. Similarly, Feldmann (1956) reported a vegetative fragmentation of sporogenic plasmodia occurring early in the growing season, during which diploid multinucleate reproductive elements (‘schizonts’) were formed. To our best knowledge, such a phenomenon has not been described in any other phytomyxid representative, and further investigation will be needed to clarify whether asexual reproduction happens at the secondary infection phase level.

As predicted by Feldmann (1956), a sexual (diploid) phase is likely initiated by conjugating secondary zoospores in *Feldmanniella* and ends with meiosis in sporogenic plasmodium prior to its cleavage into resting spores. As usual for a sexual reproduction strategy, the secondary cortical infection resulting in the formation of shoot galls is relatively rare in nature and in this case apparently linked to summer and early autumn months (Feldmann 1956, den Hartog 1989). While the processes behind this seasonality are unclear, it has now been reported in several other phytomyxid representatives such as *Tetramyxa parasitica, Marinomyxa marina* and *Ligniera verrucosa* (den Hartog 1963, Kolátková et al. 2020, Hittorf et al. 2020, Kolátková et al. 2022). For all the mentioned species, the occurrence of galls seems to somewhat correlate with the prominent blooming season of the hosts (den Hartog 1963, Hittorf et al. 2020, Kolátková et al. 2022). In *Tetramyxa parasitica*, the lower salinity of the ambient water was also suggested as one of the potential factors correlated with the formation of shoot galls (den Hartog 1963).

The secondary infection stages of *F. radicapillae* remain unknown for now. We have observed thousands of *Z. marina* shoots in nature and not once have we noticed anything resembling shoot galls. Considering the size of *Z. marina*, it is plausible that secondary plasmodia also develop within the host’s cortical tissue but do not cause the evident morphological changes typical for dwarf eelgrass species and therefore remain overlooked. Additionally, there might be lower environmental pressure on *F. radicapillae* to produce resting spores as populations of *Z. marina* are often perennial and grow in subtidal areas where the conditions are much more favourable for survival than in exposed intertidal flats where dwarf eelgrass species hosting *F. bicaudata* are found. On the other hand, life cycles of phytomyxids do not always follow the generalized scheme, and secondary infection may take place in non-cortical tissues of the host (Barr 1979, Neuhauser & Kirchmair 2009, Hittorf et al. 2022). There is a possibility that we have, for instance, simply failed to notice the later developmental stages of the *F. radicapillae* in the examined root tissue. However, we find this explanation to be rather unlikely given the number of investigated *Z. marina* specimens.

Although phytomyxids have been traditionally perceived as rare entities in seagrass meadows (den Hartog 1965, den Hartog 1989, Lipkin and Avidor 1974), Elliott et al. (2019) demonstrated at four distant geographic locations in Puget Sound, WA, USA that they are actually highly prevalent in the roots of *Z. marina* and surrounding sediments. Our present analysis of a global dataset of *Z. marina* samples (Ettinger et al. 2021) further reveals that *Feldmanniella* spp. are ubiquitous and potentially one of the most abundant protists associated with eelgrass roots on a global scale. The omnipresence of phytomyxids in eelgrass beds together with their high host-specificity indicate that they are highly specialized members of coastal/estuarine ecosystems, but most likely not substantially harmful to their hosts. Recent observation of seeds in a specimen of an invasive seagrass, *Halophila stipulacea*, heavily infested with a phytomyxid parasite *Marinomyxa marina* proves that seagrasses are still capable of sexual reproduction even when infected by these protists (Kolátková et al. 2022). We hypothesize that the severe pathogenicity attributed to some of the representatives of Plasmodiophorida parasitizing agricultural crops is an outcome of the unstable character of agrocenoses where the plants (*i*.*e*., hosts) are artificially bred and cultivated in large monocultures, and that it does not correctly reflect the role of phytomyxids in natural habitats. Nevertheless, as intracellular parasites, phytomyxids undoubtedly count as a stressor in seagrass populations and should be considered in future health assessments and ecological studies concerning seagrass stress threshold. Given the number of environmental and anthropogenic pressures currently posed on coastal ecosystems, it is not unlikely that naturally occurring diseases might eventually become one of the straws breaking the seagrass’ back.

## ACKNOWLEDGEMENTS

The authors would like to thank Galina Livingstone and Simon Phillips for their help during the search for infected specimens of *Zostera noltei* in Zandkreek, Netherlands and *Zostera japonica* in Roberts Bank, BC, respectively; Amy Replogle, Michal Morrison, and Oscar Sosa for technical assistance with sample preparation and analysis; and Brent Gowen and Louise Page for their guidance during the performed TEM analysis.

