## Supplementary figures and images for "Eelgrass (*Zostera* spp.) associated phytomyxids are host-specific congeneric parasites and predominant eukaryotes in the eelgrass rhizosphere on a global scale"

### Supplemental Figure 1

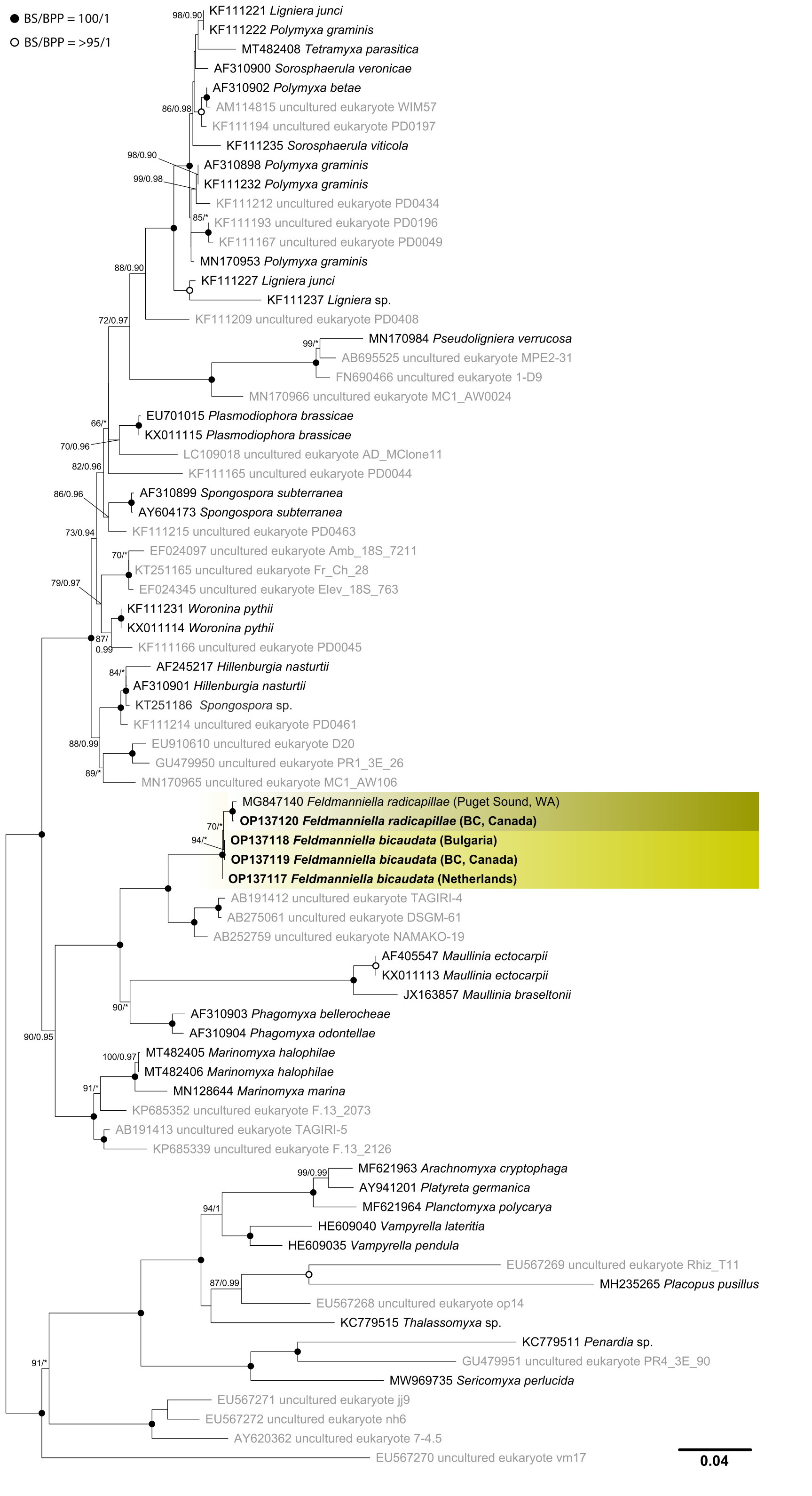
