## Supplemental Tables 1,2 for "Eelgrass (*Zostera* spp.) associated phytomyxids are host-specific congeneric parasites and predominant eukaryotes in the eelgrass rhizosphere on a global scale"

**Table S1** Summary of the proportional representation and general presence of *Feldmanniella radicapillae* OTU per investigated site (data retrieved from Ettinger et al. 2021 study)

| locality | substrate | total # of reads (*Z.marina* OTU excluded) | *Feldmanniella radicapillae* (OUT counts) | *F. radicapillae* OTU (%) | samples with *F. radicapillae* present/total # of samples |
|---|---|---|---|---|---|
| Alaska - North | leaf | 554832 | 0 | **0** | **0/11** |
| | root | 933403 | 318 | **0.034** | 6/12 |
| | sediment | 932516 | 159 | **0.017** | 2/12 |
| Alaska - South | leaf | 130882 | 2802 | **2.004** | **12/12** |
| | root | 727697 | 91135 | **12.323** | **12/12** |
| | sediment | 748051 | 5262 | **0.703** | 8/9 |
| California - North | leaf | 4698555 | 10704 | **0.228** | 24/36 |
| | root | 2453935 | 224553 | **9.151** | **25/25** |
| | sediment | 5307977 | 117398 | **2.212** | 25/29 |
| California - South | leaf | 412985 | 845 | **0.205** | 7/11 |
| | root | 573261 | 202221 | **35.276** | **12/12** |
| | sediment | 142690 | 3499 | **2.452** | 5/7 |
| Canada | leaf | 97438 | 135 | **0.139** | 3/11 |
| | root | 405061 | 61402 | **15.159** | 11/12 |
| | sediment | 853273 | 8052 | **0.944** | 8/9 |
| Croatia | leaf | 1060021 | 42 | **0.004** | 3/12 |
| | root | 243652 | 25920 | **10.638** | 4/5 |
| | sediment | 815727 | 13654 | **1.674** | 8/11 |
| French Mediterranean | leaf | 693301 | 61 | **0.009** | 5/12 |
| | root | 941128 | 689 | **0.073** | 5/7 |
| | sediment | 863098 | 107 | **0.012** | 3/9 |
| Japan - North | leaf | 276642 | 8 | **0.003** | 1/12 |
| | root | 480348 | 418 | **0.087** | 7/12 |
| | sediment | 99673 | 0 | **0** | **0/6** |
| Japan - South | leaf | 324100 | 108 | **0.033** | 6/12 |
| | root | 1041309 | 101078 | **9.707** | **12/12** |
| | sediment | 3961302 | 9023 | **0.228** | 10/11 |
| Massachusetts | leaf | 1116982 | 14 | **0.001** | 2/12 |
| | root | 482117 | 665 | **0.138** | 8/12 |
| | sediment | 90645 | 0 | **0** | **0/3** |
| North Carolina | leaf | 1787717 | 2148 | **0.120** | **12/12** |
| | root | 666124 | 214060 | **32.135** | **12/12** |
| | sediment | 1200834 | 14 | **0.001** | 1/5 |
| Norway | leaf | 260308 | 0 | **0** | **0/11** |
| | root | 361872 | 4077 | **1.127** | **3/3** |
| | sediment | 2727237 | 2301 | **0.084** | 6/11 |
| Portugal | leaf | 378396 | 9 | **0.002** | 1/12 |
| | root | 448773 | 471 | **0.105** | 3/8 |
| | sediment | 240455 | 16 | **0.007** | 1/10 |
| Sweden | leaf | 968605 | 0 | **0** | **0/12** |
| | root | 434966 | 837 | **0.192** | 3/4 |
| | sediment | 3341911 | 1419 | **0.042** | 8/12 |
| Wales | leaf | 1376415 | 47 | **0.003** | 3/12 |
| | root | 878474 | 923 | **0.105** | **6/6** |
| | sediment | 1366497 | 4935 | **0.361** | 11/12 |
| Washington | leaf | 216959 | 171 | **0.079** | 6/12 |
| | root | 578552 | 14587 | **2.521** | 11/12 |
| | sediment | 82033 | 468 | **0.571** | 5/8 |

**Table S2** Proportional representation of *Feldmanniella radicapillae* OTU in individual datasets retrieved from Ettinger et al. (2021) study

| locality (GPS) | substrate | sample ID | total # of reads | *Zostera marina* (OTU counts) | total # of reads (*Z.marina* OTU excluded) | *Feldmanniella radicapillae* (OTU counts) | *F. radicapillae* OTU (%) |
|---|---|---|---|---|---|---|---|
| Alaska - North (64.485428 N, 164.76189 W) | leaf | SRR12803376 | 103843 | 12851 | 90992 | 0 | 0 |
| | leaf | SRR12803379 | 61354 | 19383 | 41971 | 0 | 0 |
| | leaf | SRR12803443 | 93758 | 52980 | 40778 | 0 | 0 |
| | leaf | SRR12803483 | 85404 | 33850 | 51554 | 0 | 0 |
| | leaf | SRR12803545 | 62217 | 18232 | 43985 | 0 | 0 |
| | leaf | SRR12803556 | 78809 | 47768 | 31041 | 0 | 0 |
| | leaf | SRR12803557 | 81777 | 16519 | 65258 | 0 | 0 |
| | leaf | SRR12803558 | 79806 | 56071 | 23735 | 0 | 0 |
| | leaf | SRR12803699 | 85086 | 33950 | 51136 | 0 | 0 |
| | leaf | SRR12803724 | 61164 | 14810 | 46354 | 0 | 0 |
| | leaf | SRR12803962 | 92189 | 24161 | 68028 | 0 | 0 |
| | | | **885407** | **330575** | **554832** | **0** | **0** |
| Alaska - North (64.485428 N, 164.76189 W) | root | SRR12803795 | 211680 | 40817 | 170863 | 0 | 0 |
| | root | SRR12803797 | 189305 | 67518 | 121787 | 8 | 0.007 |
| | root | SRR12803798 | 236157 | 83917 | 152240 | 31 | 0.020 |
| | root | SRR12803799 | 118535 | 29621 | 88914 | 0 | 0 |
| | root | SRR12803800 | 179135 | 81432 | 97703 | 13 | 0.013 |
| | root | SRR12803801 | 210375 | 155585 | 54790 | 17 | 0.031 |
| | root | SRR12803802 | 163143 | 122703 | 40440 | 0 | 0 |
| | root | SRR12803803 | 96485 | 63755 | 32730 | 0 | 0 |
| | root | SRR12803804 | 177901 | 118788 | 59113 | 0 | 0 |
| | root | SRR12803805 | 102257 | 66984 | 35273 | 0 | 0 |
| | root | SRR12803806 | 200780 | 167985 | 32795 | 4 | 0.012 |
| | root | SRR12803808 | 139542 | 92787 | 46755 | 245 | 0.524 |
| | | | **2025295** | **1091892** | **933403** | **318** | **0.034** |
| Alaska - North (64.485428 N, 164.76189 W) | sediment | SRR12803382 | 45182 | 5768 | 39414 | 0 | 0 |
| | sediment | SRR12803383 | 73943 | 739 | 73204 | 0 | 0 |
| | sediment | SRR12803384 | 21372 | 449 | 20923 | 0 | 0 |
| | sediment | SRR12803385 | 67641 | 3133 | 64508 | 0 | 0 |
| | sediment | SRR12803386 | 24008 | 6367 | 17641 | 0 | 0 |
| | sediment | SRR12803387 | 57583 | 11340 | 46243 | 0 | 0 |
| | sediment | SRR12803388 | 131305 | 7851 | 123454 | 90 | 0.073 |
| | sediment | SRR12803389 | 222214 | 24258 | 197956 | 69 | 0.035 |
| | sediment | SRR12803391 | 124710 | 6642 | 118068 | 0 | 0 |
| | sediment | SRR12803392 | 139475 | 10479 | 128996 | 0 | 0 |
| | sediment | SRR12803393 | 81424 | 1991 | 79433 | 0 | 0 |
| | sediment | SRR12803394 | 22776 | 100 | 22676 | 0 | 0 |
| | | | **1011633** | **79117** | **932516** | **159** | **0.017** |
| Alaska - South (55.328899 N, 162.821207 W) | leaf | SRR12803568 | 49329 | 25448 | 23881 | 571 | 2.391 |
| | leaf | SRR12803579 | 37958 | 34434 | 3524 | 12 | 0.341 |
| | leaf | SRR12803590 | 19561 | 18807 | 754 | 16 | 2.122 |
| | leaf | SRR12803601 | 35203 | 23275 | 11928 | 22 | 0.184 |
| | leaf | SRR12803881 | 49699 | 17544 | 32155 | 308 | 0.958 |
| | leaf | SRR12803892 | 15210 | 14503 | 707 | 120 | 16.973 |
| | leaf | SRR12803903 | 40660 | 17727 | 22933 | 138 | 0.602 |
| | leaf | SRR12803914 | 26948 | 20297 | 6651 | 511 | 7.683 |
| | leaf | SRR12803925 | 17669 | 13345 | 4324 | 430 | 9.944 |
| | leaf | SRR12803936 | 27386 | 8415 | 18971 | 415 | 2.188 |
| | leaf | SRR12803947 | 24498 | 19444 | 5054 | 259 | 5.125 |
| | leaf | SRR12804014 | 41794 | 27267 | 14527 | 112 | 0.771 |
| | | | **385915** | **240506** | **145409** | **2914** | **2.004** |
| Alaska - South (55.328899 N, 162.821207 W) | root | SRR12803728 | 195630 | 80857 | 114773 | 9852 | 8.584 |
| | root | SRR12803729 | 9644 | 3031 | 6613 | 1428 | 21.594 |
| | root | SRR12803730 | 36850 | 5291 | 31559 | 53 | 0.168 |
| | root | SRR12803731 | 111350 | 56264 | 55086 | 23026 | 41.800 |
| | root | SRR12803732 | 172455 | 47708 | 124747 | 26716 | 21.416 |
| | root | SRR12803733 | 134481 | 67736 | 66745 | 16690 | 25.006 |
| | root | SRR12803734 | 86576 | 9497 | 77079 | 1292 | 1.676 |
| | root | SRR12803736 | 111412 | 80171 | 31241 | 5174 | 16.562 |
| | root | SRR12803737 | 144059 | 74957 | 69102 | 2580 | 3.734 |
| | root | SRR12803738 | 71598 | 7516 | 64082 | 297 | 0.463 |
| | root | SRR12803739 | 162797 | 76127 | 86670 | 4027 | 4.646 |
| | root | SRR12803740 | 144555 | 82117 | 62438 | 6232 | 9.981 |
| | | | **1381407** | **591272** | **790135** | **97367** | **12.323** |
| Alaska - South (55.328899 N, 162.821207 W) | sediment | SRR12803321 | 171827 | 39387 | 132440 | 30 | 0.023 |
| | sediment | SRR12803322 | 113019 | 47570 | 65449 | 1013 | 1.548 |
| | sediment | SRR12803323 | 82855 | 45246 | 37609 | 672 | 1.787 |
| | sediment | SRR12803324 | 65580 | 47438 | 18142 | 99 | 0.546 |
| | sediment | SRR12803325 | 132480 | 63644 | 68836 | 3113 | 4.522 |
| | sediment | SRR12803867 | 52850 | 39202 | 13648 | 297 | 2.176 |
| | sediment | SRR12803868 | 29519 | 11363 | 18156 | 7 | 0.039 |
| | sediment | SRR12803871 | 391770 | 18174 | 373596 | 31 | 0.008 |
| | sediment | SRR12803873 | 25793 | 5618 | 20175 | 0 | 0 |
| | | | **1065693** | **317642** | **748051** | **5262** | **0.703** |
| California - North (38.319755 N, 123.055136 W) | leaf | SRR12803501 | 544042 | 72542 | 471500 | 0 | 0 |
| | leaf | SRR12803502 | 451603 | 54507 | 397096 | 23 | 0.006 |
| | leaf | SRR12803503 | 408838 | 24845 | 383993 | 0 | 0 |
| | leaf | SRR12803504 | 448202 | 49149 | 399053 | 0 | 0 |
| | leaf | SRR12803513 | 139281 | 25090 | 114191 | 52 | 0.046 |
| | leaf | SRR12803514 | 181528 | 31478 | 150050 | 109 | 0.073 |
| | leaf | SRR12803515 | 135714 | 35796 | 99918 | 7 | 0.007 |
| | leaf | SRR12803516 | 189733 | 31745 | 157988 | 87 | 0.055 |
| | leaf | SRR12803517 | 121585 | 38692 | 82893 | 19 | 0.023 |
| | leaf | SRR12803518 | 102846 | 22590 | 80256 | 66 | 0.082 |
| | leaf | SRR12803519 | 6157 | 1477 | 4680 | 9 | 0.192 |
| | leaf | SRR12803520 | 215404 | 44372 | 171032 | 122 | 0.071 |
| | leaf | SRR12803521 | 168259 | 35157 | 133102 | 10 | 0.008 |
| | leaf | SRR12803522 | 208867 | 40884 | 167983 | 376 | 0.224 |
| | leaf | SRR12803524 | 68161 | 16689 | 51472 | 234 | 0.455 |
| | leaf | SRR12803525 | 109288 | 25763 | 83525 | 375 | 0.449 |
| | leaf | SRR12803526 | 159512 | 28114 | 131398 | 119 | 0.091 |
| | leaf | SRR12803527 | 157808 | 37005 | 120803 | 18 | 0.015 |
| | leaf | SRR12803528 | 154376 | 31495 | 122881 | 336 | 0.273 |
| | leaf | SRR12803529 | 138814 | 21182 | 117632 | 2333 | 1.983 |
| | leaf | SRR12803530 | 220068 | 45853 | 174215 | 354 | 0.203 |
| | leaf | SRR12803531 | 181729 | 39286 | 142443 | 227 | 0.159 |
| | leaf | SRR12803532 | 80994 | 10127 | 70867 | 0 | 0 |
| | leaf | SRR12803533 | 178485 | 66548 | 111937 | 10 | 0.009 |
| | leaf | SRR12803535 | 93124 | 11785 | 81339 | 0 | 0 |
| | leaf | SRR12803536 | 156780 | 29129 | 127651 | 23 | 0.018 |
| | leaf | SRR12803537 | 156507 | 17006 | 139501 | 172 | 0.123 |
| | leaf | SRR12803538 | 151851 | 36188 | 115663 | 5567 | 4.813 |
| | leaf | SRR12803615 | 61964 | 9135 | 52829 | 0 | 0 |
| | leaf | SRR12803626 | 48147 | 19707 | 28440 | 0 | 0 |
| | leaf | SRR12803637 | 70340 | 31178 | 39162 | 0 | 0 |
| | leaf | SRR12803698 | 45462 | 5704 | 39758 | 0 | 0 |
| | leaf | SRR12803826 | 73491 | 29163 | 44328 | 0 | 0 |
| | leaf | SRR12803837 | 27756 | 1149 | 26607 | 0 | 0 |
| | leaf | SRR12803995 | 26554 | 2839 | 23715 | 0 | 0 |
| | leaf | SRR12804006 | 50032 | 11378 | 38654 | 56 | 0.145 |

| | | | | | | | |
|---|---|---|---|---|---|---|---|
| | | | **5733302** | **1034747** | **4698555** | **10704** | **0.228** |
| California - North (38.319755 N, 123.055136 W) | root | SRR12803496 | 533162 | 317275 | 215887 | 3765 | 1.744 |
| | root | SRR12803497 | 348637 | 318073 | 30564 | 2974 | 9.730 |
| | root | SRR12803498 | 368625 | 302099 | 66526 | 1187 | 1.784 |
| | root | SRR12803500 | 395710 | 317910 | 77800 | 2306 | 2.964 |
| | root | SRR12803571 | 159416 | 85934 | 73482 | 4153 | 5.652 |
| | root | SRR12803572 | 256628 | 141191 | 115437 | 14061 | 12.181 |
| | root | SRR12803574 | 570109 | 385059 | 185050 | 1211 | 0.654 |
| | root | SRR12803577 | 531269 | 289923 | 241346 | 19326 | 8.008 |
| | root | SRR12803578 | 422703 | 272187 | 150516 | 25319 | 16.821 |
| | root | SRR12803583 | 280351 | 121464 | 158887 | 25703 | 16.177 |
| | root | SRR12803584 | 341155 | 132412 | 208743 | 17224 | 8.251 |
| | root | SRR12803586 | 291915 | 188317 | 103598 | 34474 | 33.277 |
| | root | SRR12803588 | 396638 | 269422 | 127216 | 14029 | 11.028 |
| | root | SRR12803589 | 409217 | 233367 | 175850 | 711 | 0.404 |
| | root | SRR12803593 | 126080 | 43209 | 82871 | 9426 | 11.374 |
| | root | SRR12803594 | 165308 | 103864 | 61444 | 8280 | 13.476 |
| | root | SRR12803595 | 169905 | 142879 | 27026 | 879 | 3.252 |
| | root | SRR12803677 | 298504 | 265543 | 32961 | 1516 | 4.599 |
| | root | SRR12803678 | 222959 | 148937 | 74022 | 13360 | 18.049 |
| | root | SRR12803679 | 264596 | 200381 | 64215 | 20010 | 31.161 |
| | root | SRR12803680 | 134647 | 124747 | 9900 | 666 | 6.727 |
| | root | SRR12803681 | 156154 | 114528 | 41626 | 16 | 0.038 |
| | root | SRR12803682 | 149562 | 128157 | 21405 | 329 | 1.537 |
| | root | SRR12803683 | 146431 | 103313 | 43118 | 523 | 1.213 |
| | root | SRR12803685 | 100969 | 36524 | 64445 | 3105 | 4.818 |
| | | | **7240650** | **4786715** | **2453935** | **224553** | **9.151** |
| California - North (38.319755 N, 123.055136 W) | sediment | SRR12803492 | 419674 | 121957 | 297717 | 1862 | 0.625 |
| | sediment | SRR12803493 | 342169 | 133765 | 208404 | 680 | 0.326 |
| | sediment | SRR12803494 | 341085 | 216126 | 124959 | 218 | 0.174 |
| | sediment | SRR12803495 | 364134 | 181889 | 182245 | 891 | 0.489 |
| | sediment | SRR12803778 | 14691 | 10934 | 3757 | 57 | 1.517 |
| | sediment | SRR12803779 | 21789 | 12806 | 8983 | 147 | 1.636 |
| | sediment | SRR12803820 | 178463 | 22323 | 156140 | 36 | 0.023 |
| | sediment | SRR12803823 | 34775 | 8196 | 26579 | 0 | 0 |
| | sediment | SRR12803824 | 293292 | 59241 | 234051 | 389 | 0.166 |
| | sediment | SRR12803825 | 337040 | 45544 | 291496 | 142 | 0.049 |
| | sediment | SRR12803827 | 486509 | 56757 | 429752 | 255 | 0.059 |
| | sediment | SRR12803828 | 296719 | 35620 | 261099 | 1218 | 0.466 |
| | sediment | SRR12803830 | 496700 | 48290 | 448410 | 211 | 0.047 |
| | sediment | SRR12803832 | 151467 | 17324 | 134143 | 0 | 0 |
| | sediment | SRR12803833 | 350526 | 55722 | 294804 | 284 | 0.096 |
| | sediment | SRR12803835 | 236726 | 20319 | 216407 | 64 | 0.030 |
| | sediment | SRR12803836 | 93410 | 18699 | 74711 | 33 | 0.044 |
| | sediment | SRR12803838 | 291828 | 3708 | 288120 | 0 | 0 |
| | sediment | SRR12803839 | 336016 | 49517 | 286499 | 8295 | 2.895 |
| | sediment | SRR12803841 | 382829 | 35376 | 347453 | 97092 | 27.944 |
| | sediment | SRR12803842 | 65261 | 15741 | 49520 | 139 | 0.281 |
| | sediment | SRR12803843 | 77716 | 6432 | 71284 | 0 | 0 |
| | sediment | SRR12803844 | 458707 | 64778 | 393929 | 4289 | 1.089 |
| | sediment | SRR12803845 | 338560 | 6522 | 332038 | 134 | 0.040 |
| | sediment | SRR12803846 | 31103 | 24496 | 6607 | 93 | 1.408 |
| | sediment | SRR12803847 | 69133 | 37658 | 31475 | 188 | 0.597 |
| | sediment | SRR12803849 | 49162 | 24286 | 24876 | 132 | 0.531 |
| | sediment | SRR12803850 | 122059 | 62950 | 59109 | 435 | 0.736 |
| | sediment | SRR12803852 | 39839 | 16429 | 23410 | 114 | 0.487 |
| | | | **6721382** | **1413405** | **5307977** | **117398** | **2.212** |
| California - South (32.713756 N, 117.225474 W) | leaf | SRR12803484 | 51703 | 1525 | 50178 | 0 | 0 |
| | leaf | SRR12803486 | 68675 | 7870 | 60805 | 26 | 0.043 |
| | leaf | SRR12803487 | 36706 | 5633 | 31073 | 295 | 0.949 |
| | leaf | SRR12803488 | 31612 | 452 | 31160 | 0 | 0 |
| | leaf | SRR12803489 | 37391 | 17401 | 19990 | 218 | 1.091 |
| | leaf | SRR12803490 | 50773 | 6974 | 43799 | 206 | 0.470 |
| | leaf | SRR12803491 | 42203 | 2036 | 40167 | 8 | 0.020 |
| | leaf | SRR12803499 | 37635 | 1985 | 35650 | 0 | 0 |
| | leaf | SRR12803961 | 44644 | 1659 | 42985 | 13 | 0.030 |
| | leaf | SRR12803973 | 18620 | 16939 | 1681 | 79 | 4.700 |
| | leaf | SRR12803984 | 60450 | 4953 | 55497 | 0 | 0 |
| | | | **480412** | **67427** | **412985** | **845** | **0.205** |
| California - South (32.713756 N, 117.225474 W) | root | SRR12803664 | 115503 | 68376 | 47127 | 17008 | 36.090 |
| | root | SRR12803665 | 76181 | 41578 | 34603 | 21244 | 61.394 |
| | root | SRR12803666 | 92760 | 41630 | 51130 | 18108 | 35.416 |
| | root | SRR12803667 | 127356 | 77927 | 49429 | 28322 | 57.298 |
| | root | SRR12803668 | 111428 | 61890 | 49538 | 6921 | 13.971 |
| | root | SRR12803669 | 123574 | 59074 | 64500 | 2270 | 3.519 |
| | root | SRR12803670 | 116558 | 78857 | 37701 | 20875 | 55.370 |
| | root | SRR12803671 | 100628 | 51002 | 49626 | 27977 | 56.376 |
| | root | SRR12803672 | 174893 | 128956 | 45937 | 17909 | 38.986 |
| | root | SRR12803674 | 152797 | 100189 | 52608 | 27760 | 52.768 |
| | root | SRR12803675 | 174669 | 124111 | 50558 | 859 | 1.699 |
| | root | SRR12803676 | 97296 | 56792 | 40504 | 12968 | 32.017 |
| | | | **1463643** | **890382** | **573261** | **202221** | **35.276** |
| California - South (32.713756 N, 117.225474 W) | sediment | SRR12803766 | 41870 | 5114 | 36756 | 749 | 2.038 |
| | sediment | SRR12803767 | 11389 | 1486 | 9903 | 0 | 0 |
| | sediment | SRR12803768 | 15456 | 1540 | 13916 | 802 | 5.763 |
| | sediment | SRR12803772 | 6992 | 876 | 6116 | 77 | 1.259 |
| | sediment | SRR12803775 | 15069 | 4297 | 10772 | 722 | 6.703 |
| | sediment | SRR12803776 | 27302 | 77 | 27225 | 0 | 0 |
| | sediment | SRR12803777 | 47579 | 9577 | 38002 | 1149 | 3.024 |
| | | | **165657** | **22967** | **142690** | **3499** | **2.452** |
| Canada (49.11237 N, 68.17593 W) | leaf | SRR12803441 | 19525 | 13700 | 5825 | 63 | 1.082 |
| | leaf | SRR12803442 | 22033 | 8685 | 13348 | 0 | 0 |
| | leaf | SRR12803444 | 14637 | 12343 | 2294 | 0 | 0 |
| | leaf | SRR12803445 | 12276 | 5849 | 6427 | 0 | 0 |
| | leaf | SRR12803446 | 6582 | 3883 | 2699 | 0 | 0 |
| | leaf | SRR12803447 | 12530 | 11803 | 727 | 0 | 0 |
| | leaf | SRR12803448 | 23845 | 13375 | 10470 | 45 | 0.430 |
| | leaf | SRR12803449 | 43205 | 2237 | 40968 | 0 | 0 |
| | leaf | SRR12803451 | 25848 | 17697 | 8151 | 0 | 0 |
| | leaf | SRR12803481 | 10501 | 5451 | 5050 | 27 | 0.535 |
| | leaf | SRR12803482 | 13817 | 12338 | 1479 | 0 | 0 |
| | | | **204799** | **107361** | **97438** | **135** | **0.139** |
| Canada (49.11237 N, 68.17593 W) | root | SRR12803650 | 162927 | 135018 | 27909 | 4077 | 14.608 |
| | root | SRR12803652 | 149746 | 130135 | 19611 | 22 | 0.112 |
| | root | SRR12803653 | 85114 | 75014 | 10100 | 176 | 1.743 |
| | root | SRR12803654 | 111928 | 54005 | 57923 | 126 | 0.218 |
| | root | SRR12803655 | 123777 | 70767 | 53010 | 10816 | 20.404 |
| | root | SRR12803656 | 91861 | 65881 | 25980 | 287 | 1.105 |
| | root | SRR12803657 | 65243 | 52377 | 12866 | 3289 | 25.564 |
| | root | SRR12803658 | 150298 | 119203 | 31095 | 3046 | 9.796 |
| | root | SRR12803659 | 30277 | 18364 | 11913 | 51 | 0.428 |
| | root | SRR12803660 | 135325 | 87868 | 47457 | 6851 | 14.436 |
| | root | SRR12803661 | 144694 | 68288 | 76406 | 32661 | 42.747 |

| | | | | | | | |
|---|---|---|---|---|---|---|---|
| | root | SRR12803663 | 124446 | 93655 | 30791 | 0 | 0 |
| | | | **1375636** | **970575** | **405061** | **61402** | **15.159** |
| Canada (49.11237 N, 68.17593 W) | sediment | SRR12803752 | 511015 | 40547 | 470468 | 6022 | 1.280 |
| | sediment | SRR12803753 | 42900 | 26795 | 16105 | 0 | 0 |
| | sediment | SRR12803754 | 95575 | 8738 | 86837 | 425 | 0.489 |
| | sediment | SRR12803755 | 64373 | 9484 | 54889 | 310 | 0.565 |
| | sediment | SRR12803756 | 83283 | 17962 | 65321 | 613 | 0.938 |
| | sediment | SRR12803757 | 28031 | 3197 | 24834 | 143 | 0.576 |
| | sediment | SRR12803758 | 85676 | 24308 | 61368 | 419 | 0.683 |
| | sediment | SRR12803759 | 57400 | 4483 | 52917 | 65 | 0.123 |
| | sediment | SRR12803763 | 28617 | 8083 | 20534 | 55 | 0.268 |
| | | | **996870** | **143597** | **853273** | **8052** | **0.944** |
| Croatia (44.211628 N, 15.490607 E) | leaf | SRR12803362 | 168194 | 74487 | 93707 | 15 | 0.016 |
| | leaf | SRR12803363 | 146714 | 60450 | 86264 | 0 | 0 |
| | leaf | SRR12803365 | 176246 | 62119 | 114127 | 0 | 0 |
| | leaf | SRR12803366 | 104142 | 20367 | 83775 | 14 | 0.017 |
| | leaf | SRR12803367 | 158167 | 37748 | 120419 | 0 | 0 |
| | leaf | SRR12803368 | 135429 | 66231 | 69198 | 0 | 0 |
| | leaf | SRR12803369 | 109048 | 41425 | 67623 | 0 | 0 |
| | leaf | SRR12803370 | 162432 | 60472 | 101960 | 0 | 0 |
| | leaf | SRR12803371 | 87931 | 56368 | 31563 | 0 | 0 |
| | leaf | SRR12803372 | 147435 | 34769 | 112666 | 0 | 0 |
| | leaf | SRR12803373 | 106162 | 27493 | 78669 | 0 | 0 |
| | leaf | SRR12803374 | 144256 | 44206 | 100050 | 13 | 0.013 |
| | | | **1646156** | **586135** | **1060021** | **42** | **0.004** |
| Croatia (44.211628 N, 15.490607 E) | root | SRR12803893 | 445065 | 345377 | 99688 | 15547 | 15.596 |
| | root | SRR12803897 | 194752 | 156835 | 37917 | 372 | 0.981 |
| | root | SRR12803900 | 137577 | 134208 | 3369 | 0 | 0 |
| | root | SRR12803902 | 295485 | 281097 | 14388 | 68 | 0.473 |
| | root | SRR12803904 | 424938 | 336648 | 88290 | 9933 | 11.250 |
| | | | **1497817** | **1254165** | **243652** | **25920** | **10.638** |
| Croatia (44.211628 N, 15.490607 E) | sediment | SRR12803965 | 97824 | 4617 | 93207 | 9874 | 10.594 |
| | sediment | SRR12803966 | 8325 | 486 | 7839 | 0 | 0 |
| | sediment | SRR12803967 | 19756 | 4233 | 15523 | 418 | 2.693 |
| | sediment | SRR12803968 | 66428 | 1118 | 65310 | 420 | 0.643 |
| | sediment | SRR12803970 | 195648 | 12747 | 182901 | 707 | 0.387 |
| | sediment | SRR12803971 | 14253 | 311 | 13942 | 0 | 0 |
| | sediment | SRR12803972 | 64075 | 4344 | 59731 | 1862 | 3.117 |
| | sediment | SRR12803974 | 157022 | 4206 | 152816 | 177 | 0.116 |
| | sediment | SRR12803975 | 158879 | 11270 | 147609 | 153 | 0.104 |
| | sediment | SRR12803976 | 39263 | 954 | 38309 | 0 | 0 |
| | sediment | SRR12803977 | 40314 | 1774 | 38540 | 43 | 0.112 |
| | | | **861787** | **46060** | **815727** | **13654** | **1.674** |
| French Mediterranean (43.447 N, 3.661503 E) | leaf | SRR12803405 | 92904 | 21376 | 71528 | 0 | 0 |
| | leaf | SRR12803406 | 130279 | 27925 | 102354 | 0 | 0 |
| | leaf | SRR12803407 | 110787 | 55776 | 55011 | 6 | 0.011 |
| | leaf | SRR12803408 | 109000 | 71048 | 37952 | 0 | 0 |
| | leaf | SRR12803409 | 91561 | 32673 | 58888 | 12 | 0.020 |
| | leaf | SRR12803410 | 121787 | 57738 | 64049 | 9 | 0.014 |
| | leaf | SRR12803411 | 118766 | 21781 | 96985 | 0 | 0 |
| | leaf | SRR12803412 | 93751 | 56824 | 36927 | 23 | 0.062 |
| | leaf | SRR12803413 | 64861 | 60731 | 4130 | 0 | 0 |
| | leaf | SRR12803414 | 141111 | 76085 | 65026 | 11 | 0.017 |
| | leaf | SRR12803416 | 128505 | 63392 | 65113 | 0 | 0 |
| | leaf | SRR12803417 | 89449 | 54111 | 35338 | 0 | 0 |
| | | | **1292761** | **599460** | **693301** | **61** | **0.009** |
| French Mediterranean (43.447 N, 3.661503 E) | root | SRR12803906 | 235817 | 175457 | 60360 | 0 | 0 |
| | root | SRR12803907 | 7495 | 4035 | 3460 | 3 | 0.087 |
| | root | SRR12803908 | 365572 | 171305 | 194267 | 44 | 0.023 |
| | root | SRR12803909 | 652564 | 234294 | 418270 | 29 | 0.007 |
| | root | SRR12803913 | 159162 | 115973 | 43189 | 0 | 0 |
| | root | SRR12803916 | 303970 | 100226 | 203744 | 602 | 0.295 |
| | root | SRR12803918 | 39135 | 21297 | 17838 | 11 | 0.062 |
| | | | **1763715** | **822587** | **941128** | **689** | **0.073** |
| French Mediterranean (43.447 N, 3.661503 E) | sediment | SRR12803978 | 8475 | 552 | 7923 | 0 | 0 |
| | sediment | SRR12803979 | 5786 | 355 | 5431 | 32 | 0.589 |
| | sediment | SRR12803981 | 260046 | 369 | 259677 | 0 | 0 |
| | sediment | SRR12803982 | 26989 | 150 | 26839 | 0 | 0 |
| | sediment | SRR12803983 | 419320 | 88 | 419232 | 47 | 0.011 |
| | sediment | SRR12803985 | 110545 | 381 | 110164 | 28 | 0.025 |
| | sediment | SRR12803987 | 6783 | 564 | 6219 | 0 | 0 |
| | sediment | SRR12803988 | 6252 | 182 | 6070 | 0 | 0 |
| | sediment | SRR12803990 | 21784 | 241 | 21543 | 0 | 0 |
| | | | **865980** | **2882** | **863098** | **107** | **0.012** |
| Japan - North (43.021167 N, 144.903217 E) | leaf | SRR12803314 | 55533 | 24638 | 30895 | 0 | 0 |
| | leaf | SRR12803353 | 5131 | 3133 | 1998 | 0 | 0 |
| | leaf | SRR12803364 | 72307 | 39147 | 33160 | 0 | 0 |
| | leaf | SRR12803375 | 61502 | 32681 | 28821 | 0 | 0 |
| | leaf | SRR12803415 | 42081 | 35644 | 6437 | 0 | 0 |
| | leaf | SRR12803426 | 58800 | 42837 | 15963 | 0 | 0 |
| | leaf | SRR12803455 | 64718 | 28048 | 36670 | 0 | 0 |
| | leaf | SRR12803463 | 66383 | 38762 | 27621 | 0 | 0 |
| | leaf | SRR12803474 | 34003 | 25290 | 8713 | 8 | 0.092 |
| | leaf | SRR12803512 | 80043 | 21231 | 58812 | 0 | 0 |
| | leaf | SRR12803523 | 103120 | 85741 | 17379 | 0 | 0 |
| | leaf | SRR12803534 | 56477 | 46304 | 10173 | 0 | 0 |
| | | | **700098** | **423456** | **276642** | **8** | **0.003** |
| Japan - North (43.021167 N, 144.903217 E) | root | SRR12803782 | 127407 | 60910 | 66497 | 0 | 0 |
| | root | SRR12803783 | 54553 | 47114 | 7439 | 0 | 0 |
| | root | SRR12803784 | 159976 | 40669 | 119307 | 10 | 0.008 |
| | root | SRR12803786 | 133973 | 127966 | 6007 | 7 | 0.117 |
| | root | SRR12803787 | 132389 | 128950 | 3439 | 3 | 0.087 |
| | root | SRR12803788 | 148188 | 142575 | 5613 | 29 | 0.517 |
| | root | SRR12803789 | 191080 | 131631 | 59449 | 0 | 0 |
| | root | SRR12803790 | 192448 | 182012 | 10436 | 82 | 0.786 |
| | root | SRR12803791 | 171143 | 44965 | 126178 | 88 | 0.070 |
| | root | SRR12803792 | 121073 | 112883 | 8190 | 0 | 0 |
| | root | SRR12803793 | 129447 | 122500 | 6947 | 199 | 2.865 |
| | root | SRR12803794 | 194985 | 134139 | 60846 | 0 | 0 |
| | | | **1756662** | **1276314** | **480348** | **418** | **0.087** |
| Japan - North (43.021167 N, 144.903217 E) | sediment | SRR12803340 | 29656 | 4395 | 25261 | 0 | 0 |
| | sediment | SRR12803343 | 16682 | 1572 | 15110 | 0 | 0 |
| | sediment | SRR12803345 | 13484 | 1194 | 12290 | 0 | 0 |
| | sediment | SRR12803346 | 17264 | 652 | 16612 | 0 | 0 |
| | sediment | SRR12803348 | 20895 | 3106 | 17789 | 0 | 0 |
| | sediment | SRR12803380 | 13574 | 963 | 12611 | 0 | 0 |
| | | | **111555** | **11882** | **99673** | **0** | **0** |
| Japan - South (34.297834 N, 132.91631 E) | leaf | SRR12803303 | 69187 | 6866 | 62321 | 0 | 0 |
| | leaf | SRR12803640 | 90055 | 54752 | 35303 | 22 | 0.062 |
| | leaf | SRR12803651 | 73209 | 44600 | 28609 | 13 | 0.045 |
| | leaf | SRR12803662 | 56221 | 34826 | 21395 | 3 | 0.014 |
| | leaf | SRR12803673 | 68396 | 41265 | 27131 | 23 | 0.085 |
| | leaf | SRR12803684 | 113371 | 92643 | 20728 | 43 | 0.207 |

| | | | | | | | |
|---|---|---|---|---|---|---|---|
| | leaf | SRR12803695 | 14525 | 11400 | 3125 | 0 | 0 |
| | leaf | SRR12803735 | 48406 | 17818 | 30588 | 4 | 0.013 |
| | leaf | SRR12803746 | 46073 | 32812 | 13261 | 0 | 0 |
| | leaf | SRR12803785 | 46627 | 14026 | 32601 | 0 | 0 |
| | leaf | SRR12803796 | 57923 | 33365 | 24558 | 0 | 0 |
| | leaf | SRR12803807 | 74637 | 50157 | 24480 | 0 | 0 |
| | | | **758630** | **434530** | **324100** | **108** | **0.033** |
| Japan - South (34.297834 N, 132.91631 E) | root | SRR12803741 | 132276 | 94883 | 37393 | 5947 | 15.904 |
| | root | SRR12803742 | 135439 | 24699 | 110740 | 15797 | 14.265 |
| | root | SRR12803743 | 154364 | 15428 | 138936 | 11118 | 8.002 |
| | root | SRR12803744 | 134205 | 48314 | 85891 | 5759 | 6.705 |
| | root | SRR12803745 | 96290 | 24097 | 72193 | 2668 | 3.696 |
| | root | SRR12803747 | 145190 | 70408 | 74782 | 27912 | 37.324 |
| | root | SRR12803748 | 154621 | 92377 | 62244 | 13912 | 22.351 |
| | root | SRR12803749 | 124243 | 66350 | 57893 | 12590 | 21.747 |
| | root | SRR12803750 | 100515 | 25371 | 75144 | 86 | 0.114 |
| | root | SRR12803751 | 156287 | 50551 | 105736 | 4365 | 4.128 |
| | root | SRR12803780 | 128969 | 25940 | 103029 | 361 | 0.350 |
| | root | SRR12803781 | 144544 | 27216 | 117328 | 563 | 0.480 |
| | | | **1606943** | **565634** | **1041309** | **101078** | **9.707** |
| Japan - South (34.297834 N, 132.91631 E) | sediment | SRR12803326 | 125969 | 1219 | 124750 | 222 | 0.178 |
| | sediment | SRR12803327 | 356436 | 18032 | 338404 | 3746 | 1.107 |
| | sediment | SRR12803329 | 433701 | 10886 | 422815 | 1138 | 0.269 |
| | sediment | SRR12803330 | 389726 | 2136 | 387590 | 753 | 0.194 |
| | sediment | SRR12803331 | 461784 | 1958 | 459826 | 1703 | 0.370 |
| | sediment | SRR12803332 | 551534 | 4567 | 546967 | 310 | 0.057 |
| | sediment | SRR12803333 | 322905 | 702 | 322203 | 234 | 0.073 |
| | sediment | SRR12803334 | 452760 | 2820 | 449940 | 75 | 0.017 |
| | sediment | SRR12803336 | 558292 | 23912 | 534380 | 613 | 0.115 |
| | sediment | SRR12803337 | 282467 | 2869 | 279598 | 229 | 0.082 |
| | sediment | SRR12803338 | 94954 | 125 | 94829 | 0 | 0 |
| | | | **4030528** | **69226** | **3961302** | **9023** | **0.228** |
| Massachusetts (42.42014 N, 70.91544 W) | leaf | SRR12803433 | 153118 | 19691 | 133427 | 9 | 0.007 |
| | leaf | SRR12803434 | 54354 | 40177 | 14177 | 0 | 0 |
| | leaf | SRR12803435 | 155191 | 12093 | 143098 | 0 | 0 |
| | leaf | SRR12803436 | 30350 | 13472 | 16878 | 0 | 0 |
| | leaf | SRR12803437 | 54137 | 45610 | 8527 | 0 | 0 |
| | leaf | SRR12803438 | 194285 | 12506 | 181779 | 0 | 0 |
| | leaf | SRR12803439 | 131478 | 28204 | 103274 | 0 | 0 |
| | leaf | SRR12803440 | 165339 | 10998 | 154341 | 5 | 0.003 |
| | leaf | SRR12803552 | 144920 | 9003 | 135917 | 0 | 0 |
| | leaf | SRR12803553 | 95062 | 13967 | 81095 | 0 | 0 |
| | leaf | SRR12803554 | 71231 | 60280 | 10951 | 0 | 0 |
| | leaf | SRR12803555 | 139065 | 5547 | 133518 | 0 | 0 |
| | | | **1388530** | **271548** | **1116982** | **14** | **0.001** |
| Massachusetts (42.42014 N, 70.91544 W) | root | SRR12803609 | 175077 | 104665 | 70412 | 0 | 0 |
| | root | SRR12803610 | 109841 | 86148 | 23693 | 12 | 0.051 |
| | root | SRR12803611 | 286911 | 258934 | 27977 | 11 | 0.039 |
| | root | SRR12803641 | 157428 | 97590 | 59838 | 0 | 0 |
| | root | SRR12803642 | 180258 | 122507 | 57751 | 509 | 0.881 |
| | root | SRR12803643 | 134021 | 87091 | 46930 | 0 | 0 |
| | root | SRR12803644 | 183283 | 146660 | 36623 | 13 | 0.035 |
| | root | SRR12803645 | 133419 | 114675 | 18744 | 18 | 0.096 |
| | root | SRR12803646 | 103306 | 95291 | 8015 | 13 | 0.162 |
| | root | SRR12803647 | 119023 | 101696 | 17327 | 85 | 0.491 |
| | root | SRR12803648 | 140912 | 68914 | 71998 | 4 | 0.006 |
| | root | SRR12803649 | 140509 | 97700 | 42809 | 0 | 0 |
| | | | **1863988** | **1381871** | **482117** | **665** | **0.138** |
| Massachusetts (42.42014 N, 70.91544 W) | sediment | SRR12803712 | 14132 | 412 | 13720 | 0 | 0 |
| | sediment | SRR12803719 | 73856 | 2143 | 71713 | 0 | 0 |
| | sediment | SRR12803720 | 5836 | 624 | 5212 | 0 | 0 |
| | | | **93824** | **3179** | **90645** | **0** | **0.000** |
| North Carolina (34.692458 N, 76.622589 W) | leaf | SRR12803539 | 169658 | 20988 | 148670 | 76 | 0.051 |
| | leaf | SRR12803540 | 190893 | 9687 | 181206 | 38 | 0.021 |
| | leaf | SRR12803541 | 151568 | 9354 | 142214 | 78 | 0.055 |
| | leaf | SRR12803542 | 142480 | 11702 | 130778 | 329 | 0.252 |
| | leaf | SRR12803543 | 103446 | 4477 | 98969 | 277 | 0.280 |
| | leaf | SRR12803544 | 163347 | 18388 | 144959 | 155 | 0.107 |
| | leaf | SRR12803546 | 149978 | 14381 | 135597 | 256 | 0.189 |
| | leaf | SRR12803547 | 181170 | 4935 | 176235 | 383 | 0.217 |
| | leaf | SRR12803548 | 151242 | 4268 | 146974 | 157 | 0.107 |
| | leaf | SRR12803549 | 169131 | 3088 | 166043 | 149 | 0.090 |
| | leaf | SRR12803550 | 179803 | 2638 | 177165 | 49 | 0.028 |
| | leaf | SRR12803551 | 140796 | 1889 | 138907 | 201 | 0.145 |
| | | | **1893512** | **105795** | **1787717** | **2148** | **0.120** |
| North Carolina (34.692458 N, 76.622589 W) | root | SRR12803596 | 171445 | 125315 | 46130 | 221 | 0.479 |
| | root | SRR12803597 | 158795 | 131344 | 27451 | 441 | 1.606 |
| | root | SRR12803598 | 155495 | 106258 | 49237 | 1685 | 3.422 |
| | root | SRR12803599 | 188778 | 102668 | 86110 | 5650 | 6.561 |
| | root | SRR12803600 | 149760 | 93186 | 56574 | 32549 | 57.533 |
| | root | SRR12803602 | 197590 | 165593 | 31997 | 19487 | 60.903 |
| | root | SRR12803603 | 159993 | 101125 | 58868 | 11515 | 19.561 |
| | root | SRR12803604 | 187891 | 99227 | 88664 | 66208 | 74.673 |
| | root | SRR12803605 | 149839 | 89976 | 59863 | 4118 | 6.879 |
| | root | SRR12803606 | 133424 | 92467 | 40957 | 21322 | 52.059 |
| | root | SRR12803607 | 128718 | 101291 | 27427 | 15454 | 56.346 |
| | root | SRR12803608 | 246995 | 154149 | 92846 | 35410 | 38.138 |
| | | | **2028723** | **1362599** | **666124** | **214060** | **32.135** |
| North Carolina (34.692458 N, 76.622589 W) | sediment | SRR12803701 | 252592 | 876 | 251716 | 0 | 0 |
| | sediment | SRR12803703 | 348575 | 182 | 348393 | 0 | 0 |
| | sediment | SRR12803704 | 410646 | 440 | 410206 | 0 | 0 |
| | sediment | SRR12803705 | 73918 | 109 | 73809 | 14 | 0.019 |
| | sediment | SRR12803709 | 117161 | 451 | 116710 | 0 | 0 |
| | | | **1202892** | **2058** | **1200834** | **14** | **0.001** |
| Norway (67.267997 N, 15.257219 E) | leaf | SRR12803472 | 15984 | 14838 | 1146 | 0 | 0 |
| | leaf | SRR12803473 | 100135 | 17806 | 82329 | 0 | 0 |
| | leaf | SRR12803475 | 47263 | 35206 | 12057 | 0 | 0 |
| | leaf | SRR12803477 | 89044 | 63826 | 25218 | 0 | 0 |
| | leaf | SRR12803478 | 17367 | 13065 | 4302 | 0 | 0 |
| | leaf | SRR12803479 | 95093 | 69382 | 25711 | 0 | 0 |
| | leaf | SRR12803507 | 111865 | 69727 | 42138 | 0 | 0 |
| | leaf | SRR12803508 | 85841 | 47801 | 38040 | 0 | 0 |
| | leaf | SRR12803509 | 66046 | 44000 | 22046 | 0 | 0 |
| | leaf | SRR12803510 | 23415 | 19095 | 4320 | 0 | 0 |
| | leaf | SRR12803511 | 45183 | 42182 | 3001 | 0 | 0 |
| | | | **697236** | **436928** | **260308** | **0** | **0.000** |
| Norway (67.267997 N, 15.257219 E) | root | SRR12803562 | 460065 | 324010 | 136055 | 1154 | 0.848 |
| | root | SRR12803565 | 372912 | 203957 | 168955 | 2836 | 1.679 |
| | root | SRR12803566 | 431308 | 374446 | 56862 | 87 | 0.153 |
| | | | **1264285** | **902413** | **361872** | **4077** | **1.127** |
| Norway (67.267997 N, 15.257219 E) | sediment | SRR12803629 | 611365 | 1536 | 609829 | 0 | 0 |
| | sediment | SRR12803630 | 66628 | 27661 | 38967 | 0 | 0 |
| | sediment | SRR12803631 | 185244 | 6413 | 178831 | 45 | 0.025 |

| | | | | | | | |
|---|---|---|---|---|---|---|---|
| | sediment | SRR12803632 | 509970 | 5454 | 504516 | 2079 | 0.412 |
| | sediment | SRR12803633 | 299398 | 2074 | 297324 | 14 | 0.005 |
| | sediment | SRR12803634 | 23271 | 53 | 23218 | 26 | 0.112 |
| | sediment | SRR12803636 | 331657 | 1097 | 330560 | 0 | 0 |
| | sediment | SRR12803638 | 125394 | 2050 | 123344 | 109 | 0.088 |
| | sediment | SRR12803639 | 353636 | 2685 | 350951 | 0 | 0 |
| | sediment | SRR12803818 | 133038 | 468 | 132570 | 28 | 0.021 |
| | sediment | SRR12803819 | 159212 | 22085 | 137127 | 0 | 0 |
| | | | **2798813** | **71576** | **2727237** | **2301** | **0.084** |
| Portugal (37.01427 N, 7.493273 W) | leaf | SRR12803418 | 82854 | 80730 | 2124 | 0 | 0 |
| | leaf | SRR12803419 | 119696 | 84456 | 35240 | 0 | 0 |
| | leaf | SRR12803420 | 26638 | 25794 | 844 | 0 | 0 |
| | leaf | SRR12803421 | 67966 | 67194 | 772 | 0 | 0 |
| | leaf | SRR12803422 | 139930 | 66053 | 73877 | 9 | 0.012 |
| | leaf | SRR12803423 | 88870 | 60804 | 28066 | 0 | 0 |
| | leaf | SRR12803424 | 85201 | 42140 | 43061 | 0 | 0 |
| | leaf | SRR12803425 | 129046 | 75095 | 53951 | 0 | 0 |
| | leaf | SRR12803427 | 47208 | 38001 | 9207 | 0 | 0 |
| | leaf | SRR12803428 | 102438 | 69525 | 32913 | 0 | 0 |
| | leaf | SRR12803429 | 128839 | 43094 | 85745 | 0 | 0 |
| | leaf | SRR12803430 | 70172 | 57576 | 12596 | 0 | 0 |
| | | | **1088858** | **710462** | **378396** | **9** | **0.002** |
| Portugal (37.01427 N, 7.493273 W) | root | SRR12803919 | 223273 | 133868 | 89405 | 33 | 0.037 |
| | root | SRR12803920 | 133096 | 111223 | 21873 | 0 | 0 |
| | root | SRR12803921 | 486135 | 439982 | 46153 | 0 | 0 |
| | root | SRR12803923 | 294993 | 171353 | 123640 | 0 | 0 |
| | root | SRR12803927 | 189418 | 108199 | 81219 | 211 | 0.260 |
| | root | SRR12803928 | 11849 | 8160 | 3689 | 0 | 0 |
| | root | SRR12803930 | 211949 | 154592 | 57357 | 0 | 0 |
| | root | SRR12803931 | 132000 | 106563 | 25437 | 227 | 0.892 |
| | | | **1682713** | **1233940** | **448773** | **471** | **0.105** |
| Portugal (37.01427 N, 7.493273 W) | sediment | SRR12803991 | 14346 | 4704 | 9642 | 0 | 0 |
| | sediment | SRR12803992 | 15769 | 2414 | 13355 | 0 | 0 |
| | sediment | SRR12803994 | 12643 | 4142 | 8501 | 0 | 0 |
| | sediment | SRR12803996 | 149605 | 12349 | 137256 | 16 | 0.012 |
| | sediment | SRR12803997 | 6089 | 3422 | 2667 | 0 | 0 |
| | sediment | SRR12803998 | 8202 | 4038 | 4164 | 0 | 0 |
| | sediment | SRR12803999 | 14089 | 2279 | 11810 | 0 | 0 |
| | sediment | SRR12804000 | 24228 | 6225 | 18003 | 0 | 0 |
| | sediment | SRR12804002 | 21241 | 11750 | 9491 | 0 | 0 |
| | sediment | SRR12804003 | 29335 | 3769 | 25566 | 0 | 0 |
| | | | **295547** | **55092** | **240455** | **16** | **0.007** |
| Sweden (58.3131 N, 11.5488 E) | leaf | SRR12803461 | 71544 | 6107 | 65437 | 0 | 0 |
| | leaf | SRR12803464 | 77798 | 14135 | 63663 | 0 | 0 |
| | leaf | SRR12803465 | 128271 | 19042 | 109229 | 0 | 0 |
| | leaf | SRR12803466 | 68590 | 20206 | 48384 | 0 | 0 |
| | leaf | SRR12803467 | 128991 | 10402 | 118589 | 0 | 0 |
| | leaf | SRR12803468 | 93572 | 27322 | 66250 | 0 | 0 |
| | leaf | SRR12803469 | 96671 | 7820 | 88851 | 0 | 0 |
| | leaf | SRR12803470 | 59899 | 3505 | 56394 | 0 | 0 |
| | leaf | SRR12803471 | 123649 | 4098 | 119551 | 0 | 0 |
| | leaf | SRR12803559 | 133584 | 10137 | 123447 | 0 | 0 |
| | leaf | SRR12803560 | 131181 | 5048 | 126133 | 0 | 0 |
| | leaf | SRR12803561 | 70028 | 21914 | 48114 | 0 | 0 |
| | | | **1112234** | **143629** | **968605** | **0** | **0** |
| Sweden (58.3131 N, 11.5488 E) | root | SRR12803945 | 12804 | 3527 | 9277 | 0 | 0 |
| | root | SRR12803949 | 382681 | 169233 | 213448 | 11 | 0.005 |
| | root | SRR12803952 | 211684 | 72386 | 139298 | 572 | 0.411 |
| | root | SRR12803955 | 187343 | 114400 | 72943 | 254 | 0.348 |
| | | | **794512** | **359546** | **434966** | **837** | **0.192** |
| Sweden (58.3131 N, 11.5488 E) | sediment | SRR12803616 | 444280 | 223507 | 220773 | 22 | 0.010 |
| | sediment | SRR12803617 | 143481 | 1964 | 141517 | 0 | 0 |
| | sediment | SRR12803618 | 320588 | 61470 | 259118 | 10 | 0.004 |
| | sediment | SRR12803619 | 372237 | 190881 | 181356 | 0 | 0 |
| | sediment | SRR12803620 | 204419 | 761 | 203658 | 0 | 0 |
| | sediment | SRR12803621 | 424220 | 9483 | 414737 | 775 | 0.187 |
| | sediment | SRR12803622 | 260420 | 26130 | 234290 | 280 | 0.120 |
| | sediment | SRR12803623 | 417409 | 21909 | 395500 | 173 | 0.044 |
| | sediment | SRR12803624 | 328985 | 1577 | 327408 | 92 | 0.028 |
| | sediment | SRR12803625 | 441343 | 40737 | 400606 | 62 | 0.015 |
| | sediment | SRR12803627 | 333882 | 142689 | 191193 | 0 | 0 |
| | sediment | SRR12803628 | 372732 | 977 | 371755 | 5 | 0.001 |
| | | | **4063996** | **722085** | **3341911** | **1419** | **0.042** |
| Wales (52.990731 N, 4.450321 W) | leaf | SRR12803431 | 80579 | 17342 | 63237 | 0 | 0 |
| | leaf | SRR12803432 | 106126 | 34554 | 71572 | 10 | 0.014 |
| | leaf | SRR12803450 | 131102 | 55569 | 75533 | 0 | 0 |
| | leaf | SRR12803452 | 174568 | 45345 | 129223 | 0 | 0 |
| | leaf | SRR12803453 | 125078 | 37709 | 87369 | 0 | 0 |
| | leaf | SRR12803454 | 86180 | 15382 | 70798 | 15 | 0.021 |
| | leaf | SRR12803456 | 235136 | 14142 | 220994 | 0 | 0 |
| | leaf | SRR12803457 | 85577 | 8690 | 76887 | 0 | 0 |
| | leaf | SRR12803458 | 163711 | 15077 | 148634 | 0 | 0 |
| | leaf | SRR12803459 | 138623 | 10823 | 127800 | 0 | 0 |
| | leaf | SRR12803460 | 169411 | 7657 | 161754 | 22 | 0.014 |
| | leaf | SRR12803462 | 157342 | 14728 | 142614 | 0 | 0 |
| | | | **1653433** | **277018** | **1376415** | **47** | **0.003** |
| Wales (52.990731 N, 4.450321 W) | root | SRR12803934 | 119098 | 85775 | 33323 | 54 | 0.162 |
| | root | SRR12803935 | 306097 | 96302 | 209795 | 31 | 0.015 |
| | root | SRR12803938 | 115605 | 97882 | 17723 | 8 | 0.045 |
| | root | SRR12803939 | 309150 | 245056 | 64094 | 6 | 0.009 |
| | root | SRR12803940 | 465791 | 125320 | 340471 | 809 | 0.238 |
| | root | SRR12803942 | 227763 | 14695 | 213068 | 15 | 0.007 |
| | | | **1543504** | **665030** | **878474** | **923** | **0.105** |
| Wales (52.990731 N, 4.450321 W) | sediment | SRR12803612 | 108012 | 15595 | 92417 | 572 | 0.619 |
| | sediment | SRR12803613 | 39983 | 5219 | 34764 | 487 | 1.401 |
| | sediment | SRR12803614 | 9046 | 25 | 9021 | 11 | 0.122 |
| | sediment | SRR12804004 | 393930 | 114214 | 279716 | 1033 | 0.369 |
| | sediment | SRR12804005 | 68012 | 6708 | 61304 | 402 | 0.656 |
| | sediment | SRR12804007 | 13506 | 7051 | 6455 | 0 | 0 |
| | sediment | SRR12804008 | 38030 | 1670 | 36360 | 91 | 0.250 |
| | sediment | SRR12804009 | 236480 | 28883 | 207597 | 54 | 0.026 |
| | sediment | SRR12804010 | 341643 | 10857 | 330786 | 2081 | 0.629 |
| | sediment | SRR12804011 | 243159 | 24857 | 218302 | 29 | 0.013 |
| | sediment | SRR12804012 | 74341 | 2385 | 71956 | 26 | 0.036 |
| | sediment | SRR12804013 | 21607 | 3788 | 17819 | 149 | 0.836 |
| | | | **1587749** | **221252** | **1366497** | **4935** | **0.361** |
| Washington (46.474 N, 124.028 W) | leaf | SRR12803328 | 60774 | 30426 | 30348 | 35 | 0.115 |
| | leaf | SRR12803339 | 40852 | 24374 | 16478 | 0 | 0 |
| | leaf | SRR12803378 | 51776 | 36161 | 15615 | 5 | 0.032 |
| | leaf | SRR12803390 | 67583 | 35730 | 31853 | 0 | 0 |
| | leaf | SRR12803401 | 60159 | 39014 | 21145 | 0 | 0 |
| | leaf | SRR12803710 | 20937 | 16870 | 4067 | 0 | 0 |
| | leaf | SRR12803721 | 45069 | 25655 | 19414 | 29 | 0.149 |

| | | | | | | | |
|---|---|---|---|---|---|---|---|
| | leaf | SRR12803760 | 41962 | 30220 | 11742 | 42 | 0.358 |
| | leaf | SRR12803771 | 19000 | 10572 | 8428 | 0 | 0 |
| | leaf | SRR12803848 | 50118 | 26275 | 23843 | 22 | 0.092 |
| | leaf | SRR12803859 | 48336 | 21046 | 27290 | 38 | 0.139 |
| | leaf | SRR12803870 | 51289 | 44553 | 6736 | 0 | 0 |
| | | | **557855** | **340896** | **216959** | **171** | **0.079** |
| Washington (46.474 N, 124.028 W) | root | SRR12803686 | 153635 | 117379 | 36256 | 566 | 1.561 |
| | root | SRR12803687 | 224122 | 191096 | 33026 | 4109 | 12.442 |
| | root | SRR12803688 | 178778 | 159814 | 18964 | 40 | 0.211 |
| | root | SRR12803689 | 145163 | 100447 | 44716 | 14 | 0.031 |
| | root | SRR12803690 | 134473 | 78570 | 55903 | 32 | 0.057 |
| | root | SRR12803691 | 143912 | 110611 | 33301 | 183 | 0.550 |
| | root | SRR12803692 | 160731 | 83597 | 77134 | 1277 | 1.656 |
| | root | SRR12803693 | 134330 | 49144 | 85186 | 7975 | 9.362 |
| | root | SRR12803694 | 102413 | 47168 | 55245 | 182 | 0.329 |
| | root | SRR12803725 | 162569 | 87945 | 74624 | 157 | 0.210 |
| | root | SRR12803726 | 156491 | 96376 | 60115 | 52 | 0.087 |
| | root | SRR12803727 | 106076 | 101994 | 4082 | 0 | 0 |
| | | | **1802693** | **1224141** | **578552** | **14587** | **2.521** |
| Washington (46.474 N, 124.028 W) | sediment | SRR12803854 | 12525 | 4438 | 8087 | 90 | 1.113 |
| | sediment | SRR12803855 | 5223 | 936 | 4287 | 28 | 0.653 |
| | sediment | SRR12803856 | 13153 | 6970 | 6183 | 0 | 0 |
| | sediment | SRR12803857 | 14428 | 4175 | 10253 | 0 | 0 |
| | sediment | SRR12803858 | 17601 | 4331 | 13270 | 298 | 2.246 |
| | sediment | SRR12803860 | 28823 | 2167 | 26656 | 24 | 0.090 |
| | sediment | SRR12803863 | 8963 | 2007 | 6956 | 28 | 0.403 |
| | sediment | SRR12803864 | 12046 | 5705 | 6341 | 0 | 0 |
| | | | **112762** | **30729** | **82033** | **468** | **0.571** |
